# Behavioral and dopaminergic signatures of resilience

**DOI:** 10.1101/2022.03.18.484885

**Authors:** Lindsay Willmore, Courtney Cameron, John Yang, Ilana Witten, Annegret Falkner

## Abstract

Chronic stress can have lasting adverse consequences in some individuals, yet others are resilient to the same stressor^1,2^. While previous work found differences in the intrinsic properties of mesolimbic dopamine (DA) neurons in susceptible and resilient individuals after stress was over;^3–10^ the causal links between DA activity during stress, dynamic stress-evoked behavior, and individual differences in susceptibility and resilience are not known. Here, we record behavior and neural activity in DA projections to the nucleus accumbens (NAc, signals reward^11–14^) and to the tail striatum (TS, signals threat^15–18^) during a multiday chronic social defeat paradigm and discover behavioral and neural signatures of resilience. Using supervised and unsupervised behavioral quantification, we find that resilient and susceptible individuals employ different behavioral strategies during stress. In addition, NAc-DA (but not TS-DA) activity is higher in the proximity of the aggressor in resilient mice, consistent with a greater subjective value of the aggressor. Moreover, NAc-DA tends to be elevated at the onset of fighting back in resilient mice and at the offset of attacks in susceptible mice. To test whether DA activation during defeat can generate resilience, and if its timing with respect to behavior is critical, we performed optogenetic stimulation of NAc-DA in open-loop (randomly timed) during defeat or timed to specific behaviors using real-time pose-tracking and behavioral classification. We find that both open-loop DA activation and fighting-back-timed activation promote resilience, in both cases reorganizing behavior during defeat toward resilience-associated patterns. Attack offset-timed activation promotes avoidance during defeat but does not promote susceptibility afterwards. Together, these data suggest a model whereby, during stress, DA in the NAc can increase resilience primarily by elevating the subjective value of the stressor rather than by reinforcing particular stress-responsive behaviors.

---

To detect individual differences in resilience and susceptibility to stress, we subjected male mice to a chronic defeat stress paradigm^19^ (Fig 1a). Individuals underwent 10 consecutive days of defeat by a series of novel aggressors and then were subjected to post-hoc behavioral tests to assess resilience (Extended Data Fig. 1a). In accordance with previous work, we observed that following the 10 days of defeat, individuals exhibited different levels of socially avoidant and anhedonic behavior, resulting in a spectrum of susceptibility to resilience^3^. To identify resilient and susceptible individuals following the defeat stress, we used a social interaction (SI) test, in which mice freely explored an arena with a novel aggressor behind a mesh barrier (Fig. 1b). Unstressed control mice (N=22 mice) spent 51.6 ± 9.65% (mean ± stdev) of the time in the chamber near the restrained aggressor (SI time), but stressed animals (N=32 mice) were less social and more variable (40.9 ± 16.7%, t-Test for equal means t=2.66, p=0.01, Levene test for equal variance W=5.46, p=0.02). Stressed mice were defined as “susceptible” if they had SI times less than 1 standard deviation below the mean from unstressed controls; otherwise they were considered “resilient” (Fig. 1b, Extended Data Fig. 1b). Susceptible mice also showed anhedonia-like behavior in the sucrose preference test^20^, consuming significantly less sucrose than resilient mice (Extended Data Fig. 1c-d, 1-way ANOVA F_(51,2)_=3.21, p=0.048). Additionally, resilience was associated with more weight gained during the 10 days of defeat stress (Extended Data Fig. 1e-f, Generalized Estimating Equations (GEE, linear model accounting for correlated repeated measurements from each mouse over time), N=32 mice, z=2.01, p=0.045). Stressed mice showed reduced exploration in a 2-chamber arena compared to controls, exhibiting fewer chamber crossings (Extended Data Fig. 1g, 1-way ANOVA F_(51,2)_=3.16 p=0.05) and more immobility^21^ (Extended Data Fig. 1h, 1-way ANOVA F_(51,2)_=5.77 p=0.006).

**Fig. 1.**
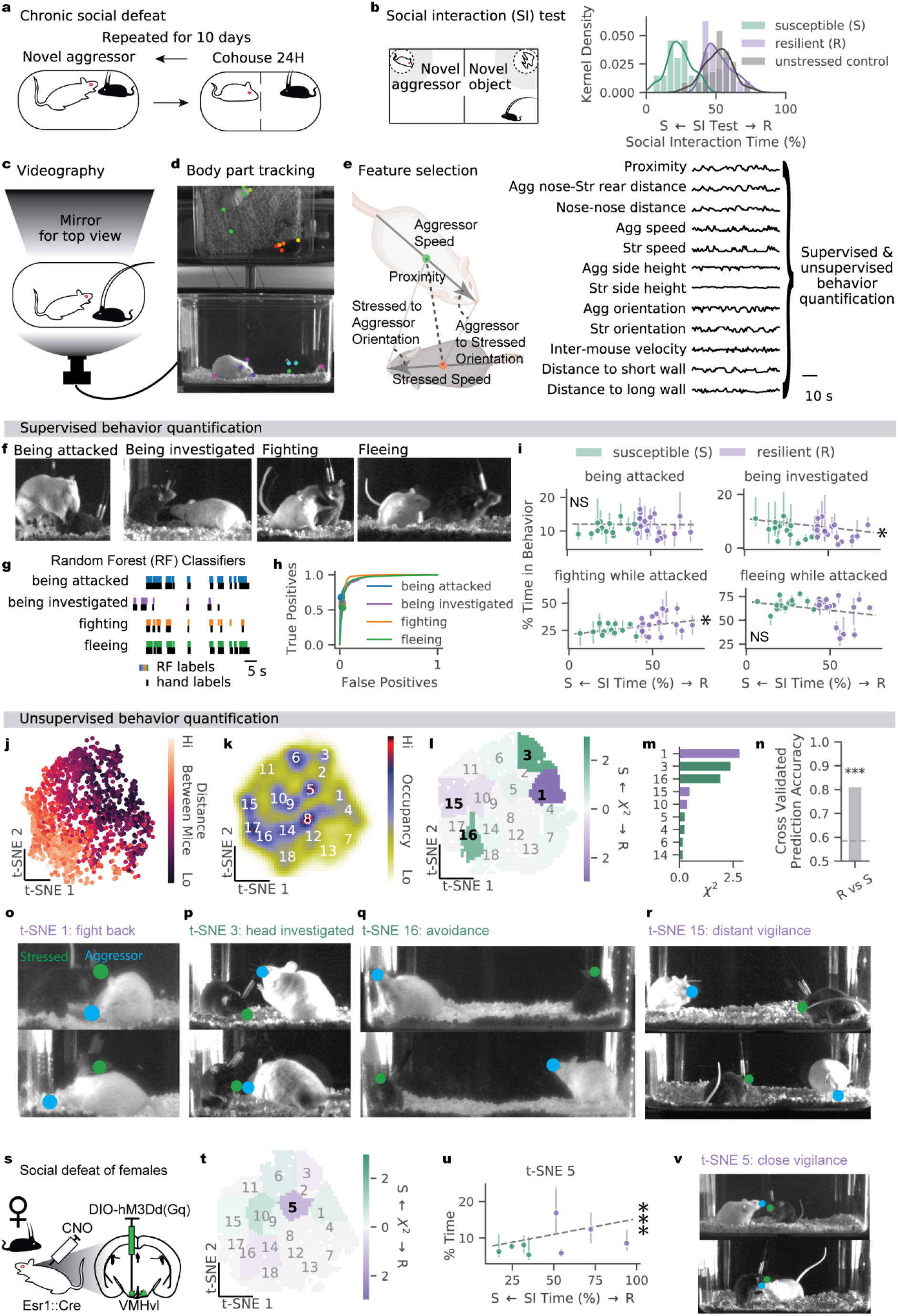
Resilience and susceptibility can be associated with different behavior profiles during defeat. **a**, Schematic of chronic social defeat stress paradigm. Interactions are recorded for quantification. **b**, Social interaction (SI) test (left). Distribution of social interaction times in unstressed control animals and stressed mice split into resilient and susceptible groups (N=22 controls, 13 susceptible, 19 resilient). **c**, Side and top-down views of defeat interactions are recorded by a single camera**. d**, Example video frame with body part tracking. **e**, Schematic of a subset of social postural features (left). Full set of 12 features used for both supervised (**f**-**i**) and unsupervised (**j**-**r**) behavior quantification (right). **f**, Example video frames of behaviors of interest. **g**, Example trace of hand annotated behavior bouts (black) and binary random forest classification (colored) of those behaviors. **h**, Receiver operator curves for behavior classification accuracy on a validation set of video frames. **i**, Relationship between time spent by individual mice in each behavior of interest and SI time (GEE: being attacked p=0.909; being investigated p=0.040; fighting while attacked, p=0.016; fleeing while attacked p=0.104; N=32 mice; means plotted with 30th to 70th percentile error bands across the 10 defeat days; N.S., not significant). **j**, t-SNE embedding from an example defeat session colored by distance between mice. **k**, Smoothed histogram of all stressed mouse behavior with behavior clusters numbered in order of increasing distance between mice. **l,** Difference in occupancy of t-SNE clusters between resilient and susceptible mice, measured by individual clusters’ chi^2^ statistics; bold numbers label 4 behaviors with greatest occupancy difference (shown in **o**-**r**). **m**, Behaviors with the largest chi^2^ statistics differentiating resilient and susceptible mice (purple bars indicate resilient-biased behaviors, green indicate susceptible-biased behaviors). **n**, Leave one mouse out cross validation accuracy (0.81) of predicting resilience vs susceptibility from an individual’s average occupancy of the t-SNE-based behavior clusters, dotted line shows 1 SD above change (0.5) accuracy with shuffled data (one-sided normal test for proportion > 0.5, p<0.001). **o-r**, Two example video frames from the 4 behavior clusters with the largest chi^2^ statistics differentiating resilient and susceptible mice (bold in **l** and top 4 shown in **m**). **s**, Schematic of chronic defeat stress with females and aggression induced via expression of excitatory DREADDs in the ventromedial hypothalamus ventrolateral area (VMHvl) of male Esr1::Cre aggressors. **t**, Difference in occupancy of t-SNE clusters between resilient (N=4) and susceptible (N=4) females, measured by individual clusters’ chi^2^ statistics; bold 5, cluster with greatest difference. **u**, Relationship between time spent by individual mice in cluster 5 and SI time (GEE: Z=4.58, p<0.001; means plotted with 30th to 70th percentile error bands across the 10 defeat days). **v**, Two example video frames from cluster 5: “close vigilance.”

While previous studies have found resilience to be predicted by behavioral and physiological measurements taken prior to defeat^22–26^, in our data, animals’ pre-defeat weights, social rank, and time spent freely interacting with conspecifics did not predict susceptibility and resilience (Extended Data Fig. 1e,i-n).

Since social behaviors can be highly variable across individuals, we hypothesized that the specific behaviors mice engaged in during defeat may predict resilience. Previously, human labeling has revealed that susceptible animals more quickly adopt submissive postures or flee more than resilient ones^27,28^. To automate behavioral quantification, we recorded every defeat session via 120 Hz videography with both top and side acquisition (Fig. 1c). Body parts of the interacting mice were tracked using markerless pose estimation software^29^ (Fig. 1d). From these body part locations, we defined 12 features to describe the animals’ relative and global postures (Fig. 1e) in order to identify four behaviors of interest: [1] being attacked, [2] being investigated, [3] fighting back, and [4] fleeing (Fig. 1f). To automate behavior identification across our large video dataset (~14 million frames), we trained binary random forest classifiers to use these features to predict human labeling of each of these behaviors (Fig. 1g,h for performance on held-out videos). Resilient and susceptible mice experienced similar levels of attack, suggesting that resilient animals do not undergo a less stressful experience (Fig. 1i). However, we identified modest behavioral differences in two other classified behaviors: 1) susceptibility was associated with more investigation from the aggressor (Fig. 1i, top, GEE, N=32 mice, z=-2.05, p=0.040), and 2) resilience was related to more often fighting back while attacked (Fig. 1i, GEE, N=32 mice, z=2.42, p=0.016, Extended Data Fig. 2 for longitudinal changes in behavior).

To complement supervised classification and to describe behavior in a more holistic manner, we used unsupervised clustering to generate behavior maps. To do this, we embedded the same 12 features from each video frame (Fig. 1e) into a 2D t-SNE manifold (t-distributed stochastic neighbor embedding; a non-linear dimensionality reduction technique to preserve local similarity and clusters in data). The embedding retained some structure from the original feature space, such as the distance between animal centroids (Fig. 1j, Extended Data Fig. 3a). We discretized behavior space into 18 clusters representing areas of peak occupancy, which, for interpretability, we numbered from those in which mice were closest together (1) to furthest apart (18) (Fig. 1k). While differences in individual cluster occupancies between susceptible and resilient individuals were small, individuals could be classified into resilient or susceptible groups well above chance accuracy using the set of all 18 cluster occupancies, suggesting differences in overall behavioral strategy (Fig. 1l-n, Extended Data Fig. 3b, 81.25% accuracy on correct classification in leave-one-mouse-out cross validation; for each mouse, a model was trained on all other mice and tested on the held out mouse). Descriptions of the behavior in a t-SNE cluster may be derived from the distribution of original features in the cluster (Extended Data Fig. 3a), the density of expression of random-forest-detected behaviors in the cluster (Extended Data Fig. 3c), and example video frames from the cluster. The 4 most informative clusters in differentiating susceptible and resilient groups included a proximal and distant behavior more often expressed by each group (Fig. 1l,m). The most resilience-associated proximal cluster (1) overlapped with fighting back frames (Fig. 1o, Extended Data Fig. 3c). The most susceptibility-associated proximal cluster (3) (Fig. 1p) can be characterized by being investigated in the head and neural implant, consistent with passive freezing^30^ and receiving orofacial sniffing^31,32^ (related to social subordination). The most susceptibility-associated distant cluster was cluster (16) “avoidance,” facing away from the aggressor (Fig. 1q). The most resilience-associated distant cluster (15) was a posture of distant “vigilance”^33^ (Fig. 1r) (Extended Data Fig. 3a).

Females, like males, have also been shown to exhibit social avoidance and anxiety-like behaviors consistent with resilience and susceptibility following stress^34–36^, though much less is known about the drivers of individual variability in females. As the behaviors that we identified in males as being resilience-associated may be sex-specific defensive strategies, we exposed females to 10 days of defeat and explored if they also have behavior patterns associated with resilience. To produce aggression towards females, we virally expressed a Cre-dependent excitatory DREADD in the ventromedial hypothalamus ventrolateral area (VMHvl) of Esr1::Cre aggressor males, and these animals were injected daily with CNO before defeat^35,37^ (Fig. 1s). As in males, we identified resilient and susceptible females on the basis of post-hoc SI time relative to an unstressed control group and performed additional post-hoc testing (Extended Data Fig. 4a-e). Expression of supervised behaviors (being attacked, being investigated, fighting, and fleeing) all did not differ between resilient and susceptible females (Extended Data Fig. 4f), and female animals exhibited significantly less fighting back behavior compared to males (Extended Data Fig. 4g, t=-7.45, p<0.001), indicating the relationship between this behavioral response strategy and resilience may be male specific. However, unsupervised analysis on female behavior during defeat revealed a single behavioral cluster that was significantly correlated with resilience (cluster 5, Fig. 1t,u, GEE, N=8 mice, z=4.58, p<0.001, Bonferroni correction for comparisons across 18 clusters). In this cluster (“close vigilance”), mice were in moderate proximity to and facing the aggressor (Fig. 1v, Extended Data Fig. 3a). Together, these data uncovered a new posture associated with resilience and suggest that attack-response behaviors (fighting vs fleeing) are not associated with resilience in females.

We next tested whether individual differences in neural dynamics during defeat and during these specific behaviors would also differentiate resilient and susceptible individuals. Differences in spontaneous activity of dopaminergic (DA) neurons recorded after the 10 days of defeat stress have been shown to differ between resilient and susceptible individuals^3,5,6^, but DA activity during defeat stress itself has not been described. In addition, different DA projections in the striatum are known to perform distinct roles^38^. DA projections to the tail of the striatum (TS) are primarily from the lateral substantia nigra and are active in response to threatening^15–18^ or alarming^39^ stimuli. DA projections from the ventral tegmental area (VTA) to the nucleus accumbens (NAc) respond vigorously to rewards such as palatable food^11–14^ and non-aggressive social investigation^40,14^.

To target these areas, we implanted optical fibers in the NAc and TS of males and females genetically expressing GCaMP6f in midbrain DAT neurons^12^ and used fiber photometry^40^ to measure bulk calcium fluorescence as a proxy for neural activity and DA release (Fig. 2a, N=19 males, 8 females; in Extended Data Fig. 5 simultaneous fast scan cyclic voltammetry and fiber photometry recordings in the NAc^13^ and TS show DA concentration and fiber photometry measurements are highly related). We confirmed that TS(DAT::GCaMP6f) responds vigorously to threatening air puffs (delivered to the nose of mice while headfixed, Fig. 2b, N=19 males, 8 females, t=5.11, p<0.001), and that NAc(DAT::GCaMP6f) responds to the approach of rewarding food (Fig. 2c, N=19 males, 8 females, t=6.59, p<0.001).

**Fig. 2.**
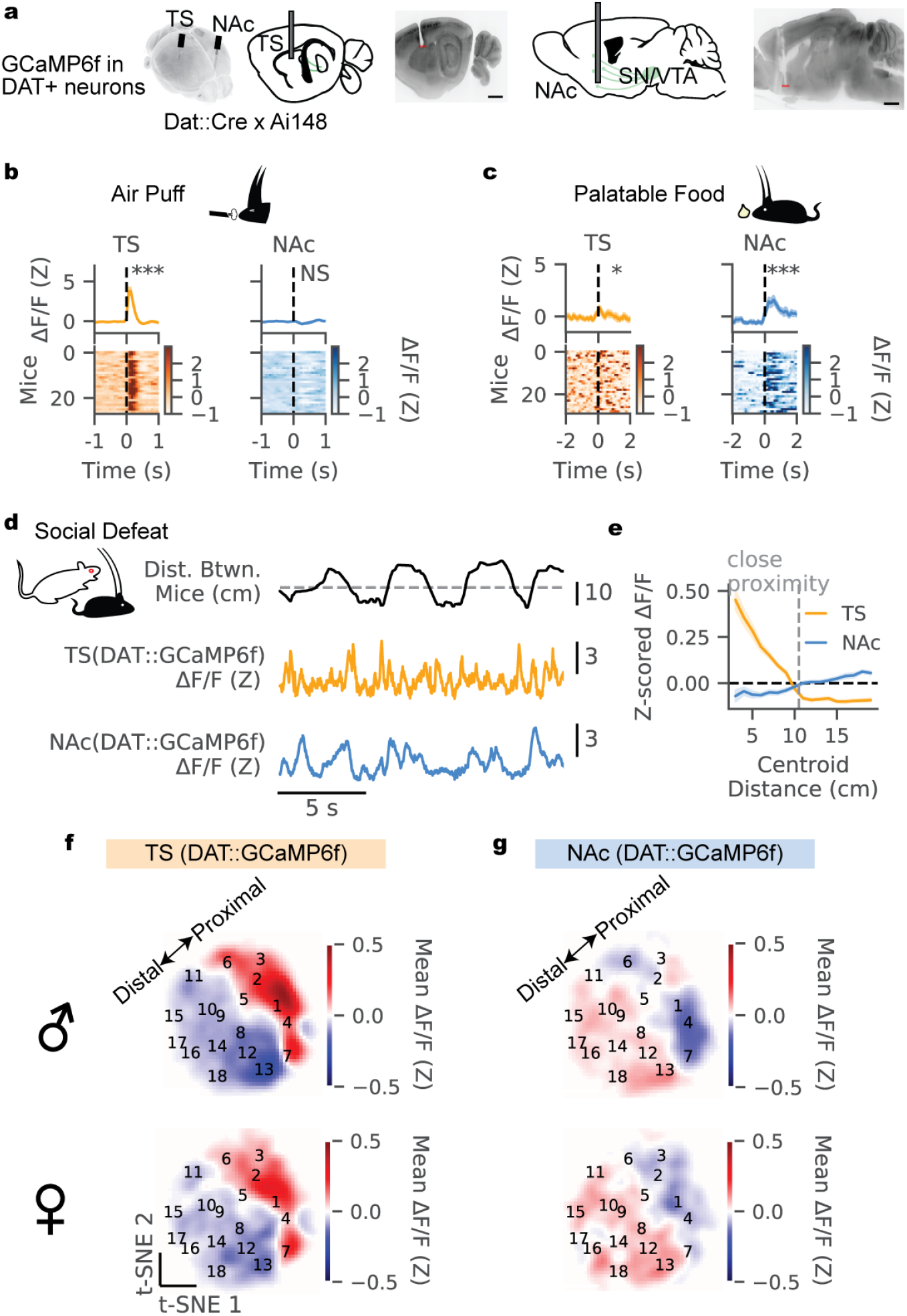
During defeat, DAT::GCaMP6f activity in the TS and NAc is oppositely modulated in proximity to the aggressor. **a,** Location of fiber photometry recordings from dopaminergic (DAT) projections in the nucleus accumbens (NAc) and tail of the striatum (TS) in the same mice, recovered using light-sheet microscopy. Scale bars (black), 1 mm. Fiber tips (red), 400 μm diameter. **b**, DAT::GCaMP6f response to air puff. Animals were head-fixed in a sliding burrow and air puffs were delivered to the snout. Average TS(DAT::GCaMP6f) and NAc(DAT::GCaMP6f) activity aligned to air puff onset within and across all mice (N=19 males, 8 females, paired t-Test for average activity after vs before puff: TS t=5.11, p<0.001, NAc t=-1.34, p=0.19)**. c**, DAT::GCaMP6f response to palatable food reward. Mice freely approach a yogurt treat dropped into their homecage. Average TS(DAT::GCaMP6f) and NAc(DAT::GCaMP6f) activity aligned to approach onset of treat within and across mice (N=19 males, 7 females, paired t-Test for average activity after vs before approach: TS t=2.36, p=0.025, NAc t=6.59, p<0.001). **d**, Example trace DAT::GCaMP6f activity in TS and NAc aligned to distance between aggressor and stressed mice with gray dotted line showing the 10.5 cm threshold used to define close proximity. **e**, Relationship between distance between mouse centroids and TS(DAT::GCaMP6f) or NAc(DAT::GCaMP6f) activity. Close proximity is defined as distances below which NAc(DAT::GCaMP6f) activity is negative (10.5 cm). **f**, Mean TS(DAT::GCaMP6f) activity across t-SNE behavior space in defeated males (top) and females (bottom). The proximal-distal line shows the axis in the behavior map defined by proximity between mice during defeat. **g**, Same as **f** for NAc(DAT::GCaMP6f).

We next explored how these DA projections, which facilitate learning stimulus-outcome associations^41,42^ and motivated actions^43,44^, may encode the defeat experience differently in resilient and susceptible mice. Recordings were performed from both sites on each day of defeat and during post-hoc testing.

First, we examined the relationship between DA activity in the TS and NAc and physical distance to the aggressor (Fig. 2d). We found that in both males and females, TS(DAT::GCaMP6f) and NAc(DAT::GCaMP6f) projections exhibited opposite profiles in response to aggressor distances; DAT::GCaMP6f activity was elevated during proximal behaviors in the TS and distal behaviors in the NAc in both male and female mice. (Fig. 2e-g). TS(DAT::GCaMP6f) activity increased at the onset of proximity to the aggressor (10.5 cm between-centroid distance) and decreased at the offset (Fig. 3a; Extended Data Fig. 6a), consistent with the threatening nature of aggressor interactions. Across the days of defeat, TS(DAT::GCaMP6f) activity at proximity onset significantly increased, indicating this threat response is experience-dependent (Fig. 2b, GEE regression for onset activity: main effect of day N=27 mice, Z=2.58, p=0.01; Extended Data Fig. 6b). However, the magnitude of TS(DAT::GCaMP6f) responses at proximity onset and offset were similar between resilient and susceptible individuals (Fig. 3b,c).

**Fig. 3.**
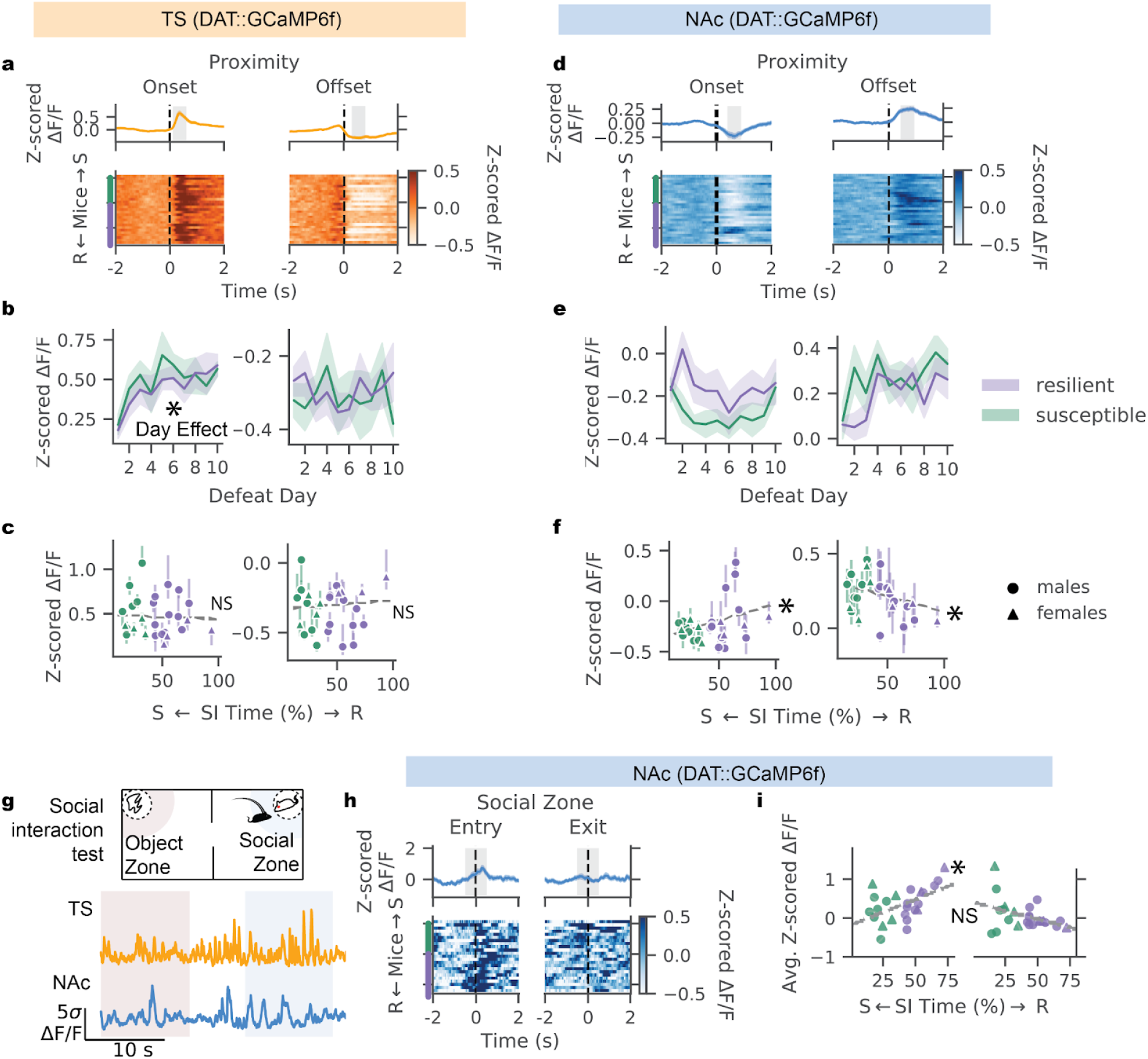
DAT::GCaMP6f in proximity to aggressors in the NAc, but not the TS, is correlated with resilience. **a**, Average aggressor proximity onset- and offset-aligned TS(DAT::GCaMP6f) activity recorded during the 10 days of defeat; average across stressed animals (N=8 females, 19 males), heatmap shows average within each mouse in order from most susceptible (green bar, N=7 males, 4 females) to most resilient (purple bar, N=12 males, 4 females). Gray region indicates 0.5s surrounding the maximum or minimum average activity following the onset or offset of proximity. **b**, Average TS(DAT::GCaMP6f) activity (in gray shaded zones in **a**) in individuals from resilient and susceptible groups across the 10 days of defeat. Solid line shows mean across mice and shaded region shows standard error. **c**, Relationship between average TS(DAT::GCaMP6f) activity (gray shaded zones in **a**) and SI time across defeated individuals (GEE regression onset activity by SI time, day and their interaction: main effect of SI time Z=0.002, p=0.6 main effect of day Z=2.584, p=0.01; GEE regression offset activity by SI time, day and their interaction: main effect of SI time Z=-0.032, p=0.98 main effect of day Z=-0.58, p=0.6). **d**, Same as **a** for NAc(DAT::GCaMP6f). **e**, Same as **b** for NAc(DAT::GCaMP6f). **f**, Same as **c** for NAc(DAT::GCaMP6f). (GEE regression onset activity by SI time, day and their interaction: main effect of SI time Z=1.98, p=0.047 main effect of day Z=-0.1, p=0.9; GEE regression offset activity by SI time, day and their interaction: main effect of SI time Z=-2.25, p=0.025 main effect of day Z=0.316, p=0.75). **g**, Schematic of fiber photometry recordings during the social interaction test, with example traces of DAT::GCaMP6f activity in the TS and NAc. **h**, Average social interaction zone entry- and exit-aligned NAc(DAT::GCaMP6f) activity across stressed animals (mean plotted with standard error, N=6 susceptible males, 4 susceptible females, 10 resilient males, 3 resilient females). **i**, Relationship between individuals’ SI time and average TS(DAT::GCaMP6f) activity in the 1s surrounding entry or exit from the social interaction zone (entry: Pearson correlation, R=0.51 p=0.01; exit: Pearson correlation, R=-0.39, p=0.062).

In contrast to the TS, we observed that NAc(DAT::GCaMP6f) activity decreased at proximity onsets and increased at offsets (Fig. 3d; Extended Data Fig. 6c). This is consistent with aggressor proximity being aversive and the end of proximal encounters providing positive “relief”^45–49^. The magnitudes of decrease at proximity onset and increase at the offset were related to susceptibility, suggesting that susceptible individuals experience more aversion to aggressor encounters and relief when encounters ended (Fig. 3e,f; GEE, N=27 mice, onset z=1.98, p=0.047; offset z=-2.25, p=0.025; Extended Data Fig. 6d).

Individual differences in DAT::GCaMP6f activity in the NAc, but not the TS, in proximity to an aggressor persisted even after the 10 days of defeat, during the post-hoc SI test (Fig. 3g-i, Extended Data Fig. 7a,b). Susceptibility was associated with less NAc(DAT::GCaMP6f) activity in entering the social zone (Fig. 3i, Pearson’s correlation; N=23 mice, 16 males, 7 females, R=0.51, p=0.01). Neither NAc(DAT::GCaMP6f) nor TS(DAT::GCaMP6f) activity during a control behavior, proximity to a novel object during the SI test, was predictive of resilience (Extended Data Fig. 7c-f). Overall, these data suggest that across both males and females, DA projections to the NAc reflect reduced subjective value of the aggressor in susceptible mice and higher aggressor value in resilient ones.

However, individual differences in NAc DA may contribute to resilience and susceptibility not only by reflecting differences in the value of social proximity, but also by correlating with - and potentially reinforcing - specific actions that may occur during proximity (e.g. fight back, flee). Potential action-specific correlates could contribute to the differences in attack-response-actions observed in resilient versus susceptible males (Fig. 1i). To test this hypothesis, we quantified the dynamics of NAc(DAT::GCaMP6f) relative to the onsets and offsets of interpretable behaviors identified using our random forest classifiers (being attacked, being investigated, fighting back, and fleeing; Fig. 1f). Since defeat-related behaviors can be close in time (for example, frames classified as being attacked and fleeing can be concurrent), behavior-triggered activity may be difficult to interpret. To account for this and to better approximate the temporal dynamics with respect to behaviors of interest, we fit the neural activity of each animal with a linear encoding model, resulting in a response profile (kernel) for each behavior^12,50^. The linear model was fit such that behavior-triggered kernels summed to best approximate the observed neural activity in each mouse (Fig. 4a-b).

**Fig. 4.**
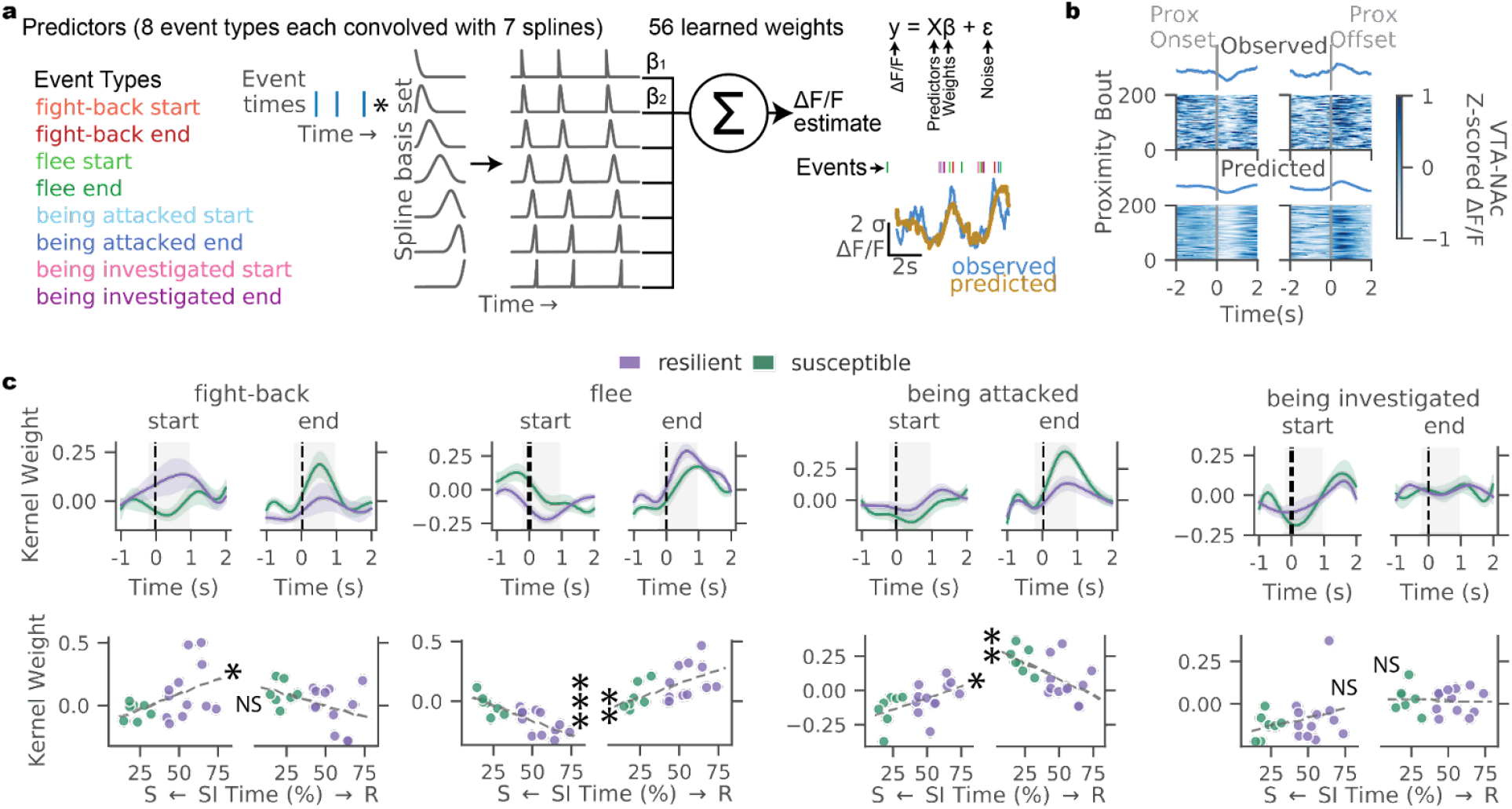
Increased NAc(DAT::GCaMP6f) at the onset fighting back is correlated with resilience while more activity at fleeing onset and attack offset are correlated with susceptibility. **a**, Schematic of the linear encoding model for the relationship between behavioral events and NAc(DAT::GCaMP6f) activity. Far left shows the full list of behavioral event types, center depicts the convolution of events of one type with 7 cubic splines spanning from 1s before to 2s after the event, and the right represents how the learned spline weights (βs) multiplied by the corresponding kernels at each event time sum to predict the neural activity. Example of behavior events and observed and predicted NAc(DAT::GCaMP6f). **b**, Example of observed (top) and model-predicted (bottom) NAc(DAT::GCaMP6f) activity across all the social proximity bouts across defeat from an example mouse, aligned to onset (left) and offset (right) of proximity (prox). **c**, Kernel weights for behavioral events, averaged across individuals in resilient and susceptible groups (top, mean and standard error plotted). Relationship between individuals’ SI time and average kernel weight in the shaded region in the above kernel weight plots (bottom, Pearson’s correlation between SI time and average kernel weight, *p<0.05, **p<0.01, ***p<0.001). In all panels, N=7 susceptible, 12 resilient mice.

Using this model, we found that resilience was associated with increased NAc(DAT::GCaMP6f) activity at the onset of fighting and decreased activity at the onset of fleeing (Fig. 4c, Pearson’s correlation, N=19 mice; R=0.49, p=0.035; R=-0.8, p=4E-5). This result is consistent with a positive relationship between resilience and fighting back behavior in males (Fig. 1i). In further support of this, males with elevated fight-related activity more often fought back (Extended Data Fig. 8a; Pearson’s correlation, N=19 mice, R=0.61, p=0.005). In contrast, susceptibility was associated with less activity at attack onset and more activity at attack offset (Fig. 4c, Pearson’s correlation, N=19 mice, R=0.47, p=0.041; R=-0.56, p=0.013), again supporting a greater aversion to attack starts and relief at attack ends in susceptible mice.

Anatomical recording location was not correlated with either kernel weights or resilience in either recording location (Extended Data Fig. 8b-d). In contrast to NAc(DAT::GCaMP6f), TS(DAT::GCaMP6f) kernels did not differ in resilient versus susceptible males (Extended Data Fig. 8e).

Overall, our recordings during defeat stress reveal that the pattern of NAc(DAT::GCaMP6f) activity is predictive of individual differences in susceptibility and resilience, and suggest at least two potential hypotheses for a causal role of these signals in conferring resilience. One hypothesis is that elevated activity in this projection during defeat could increase aggressor value, thereby producing resilience (Fig. 5a, left). This is consistent with the known role of this projection in providing a positive teaching signal for learning stimulus values^11,41,51^ and our observations that resilience was associated with greater NAc(DAT::GCaMP6f) activity at aggressor proximity both during defeat and the SI test. This hypothesis is not trivial, given previous activation of this system in the sensory period (immediately after defeat) has been shown to bias towards susceptibility rather than resilience^4^.

**Fig. 5.**
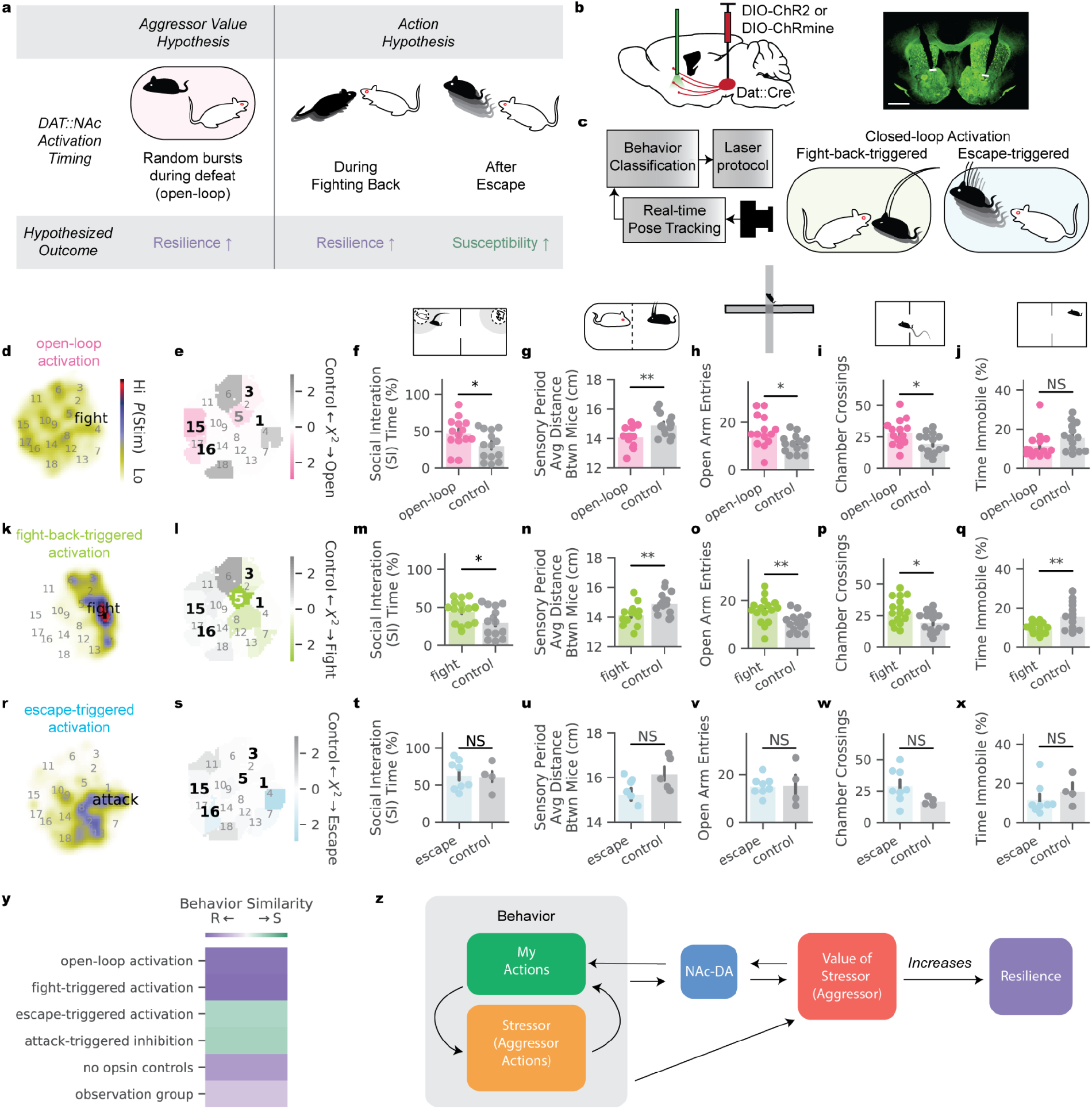
DAT::NAc activation during defeat can bias individuals towards resilience. **a**, Potential causal roles of dopaminergic (DAT) neural activity in NAc projections in generating individual differences in resilience. **b**, Schematic of neural activation strategy (left) and example histological image of optical fiber placement (right, 200 μM fiber tips in white, 1mm scale bar). **c**, Schematic of closed-loop optogenetic manipulation. **d**, Density map of where in t-SNE behavior space animals were when receiving open-loop activation. **e**, Difference in occupancy of t-SNE clusters between open-loop (N=14) and no opsin control (N=14) groups, measured by individual clusters’ chi^2^ statistics; bold numbers label behaviors differentially expressed by resilient and susceptible mice in our observational study (see Fig. 1). **f**, Social interaction (SI) time in the SI test (t-Test, difference in SI time between open-loop and no opsin control groups: t=2.19, p=0.037). **g**, Average distance between stressed and aggressor mice during the first 5 minutes of the barrier-separated sensory period across the 10 days of defeat (t-Test, difference in mean distance between open-loop and no opsin control groups: t=-2.90, p=0.008). **h**, Number of entries into the open arms of an elevated plus maze (t-Test, difference in mean entries between open-loop and no opsin control: t=2.56, p=0.016). **i**, Number of crossings between chambers of 2-chamber arena (t-Test, difference in mean crossings between open-loop and no opsin control groups: t=2.45, p=0.02). **j**, Percent of time immobile (speed < 1cm/s) during exploration of a 2-chamber arena (t-Test, difference in mean immobility between open-loop and no opsin control groups: t=-1.57, p=0.12). **k-l**, Same as **d-e** for fight-triggered activation (N=16) and no opsin controls (N=14). **m**, Same as **f** for fight-triggered activation (t-Test, t=2.29, p=0.030). **n**, Same as **g** for fight-back-triggered and no opsin controls (t-Test, t=-2.99, p=0.006). **o**, Same as **h** for fight-back-triggered and no opsin controls (t-Test, t=2.87, p=0.008). **p**, Same as **i** for fight-back-triggered and no opsin controls (t-Test, t=2.64, p=0.013). **q**, Same as **j** for fight-back-triggered and no opsin controls (t-Test, t=-3.05, p=0.005). **r-s**, Same as **d-e** for escape-triggered (N=8) and no opsin control groups (N=5). **t**, Same as **f** for escape-triggered and no opsin controls (t-Test, t=0.17, p=0.87). **u**, Same as **g** for escape-triggered activation and no opsin controls (t-Test, t=-3.94, p=0.07). **v**, Same as **h** for escape-triggered and no opsin controls (t-Test,t=-0.10, p=0.92). **w**, Same as **i** for escape-triggered and no opsin controls (t-Test, t=1.76, p=0.11). **x**, Same as **j** for escape-triggered and no opsin controls (t-Test, t=-0.80, p=0.39). **y**, Summary of relative similarity in behavior of males in manipulation and observation groups to susceptible or resilient populations from our male observational study (see Fig. 1). **z**, Model of the relationship between behaviors, dopaminergic neural activity, and resilience. We propose stimulation biased individuals towards resilience by increasing the value of the aggressor or source of stress.

Alternatively, DA may instead serve to reinforce specific actions in response to the aggressor, which may in turn impact resilience. Resilience was associated with more fighting back behavior in males (Fig. 1i) as well as more NAc(DAT::GCaMP6f) at the onset of this behavior (Fig. 4c), while susceptibility was related to reduced NAc(DAT::GCaMP6f) at the onset of this behavior and elevated NAc(DAT::GCaMP6f) at the onset of fleeing (Fig. 4c), a susceptibility-biased behavior^28^. If these differences are causally related to resilience, increasing DA during resilience-associated fighting would lead to resilience while increasing DA after susceptibility-associated fleeing would lead to susceptibility (Fig. 5a, right).

To differentiate between these two hypotheses, we performed open- and closed-loop optogenetic activation across the 10 days of social defeat followed by post-tests, including tests of anxiety and mobility. To target DA circuits, Dat::cre animals were bilaterally injected with an excitatory opsin and optical fibers were implanted in the NAc (Fig. 5b, N=38). After recovery and before defeat, opsin expression was confirmed behaviorally using a real-time place preference test (Extended Data Fig. 9a). Control animals were implanted with optical fibers but without opsin (N=19). To perform closed-loop stimulation, we developed a system for behavior-triggered optogenetic activation, using fast, online pose-detection. We streamed video frames during defeat directly to a trained pose estimation network, extracted social postural features as previously described (Fig. 1e), and applied a trained binary random forest classifier for our behavior of interest (Fig. 5c). Phasic light stimulation was triggered contingent on behavior detection.

To first test if DAT::NAc activation can produce resilience by increasing aggressor value without reinforcing specific actions, we performed open-loop stimulation during defeat, untimed to behavior. Phasic bursts of light delivery were randomly distributed throughout behavior space (Fig. 5d, burst stimulation for 2.99% of the defeat period on average, Extended Data. Fig. 9d). Animals receiving activation had greater SI times after defeat (Fig. 5f) and had reduced average distance from the aggressor during the sensory period immediately after each defeat (Fig. 5g). Furthermore, the open-loop stimulation resulted in decreased anxiety in non-social contexts: more open arm entries in the elevated plus maze (Fig. 5h) and a greater number of crossings in 2-chamber exploration (Fig. 5i). Thus, open-loop DAT::NAc activation, which was not timed to any specific action, increased appetitive social behaviors (indicative of higher aggressor value) and increased general resilience following defeat.

To directly test our second hypothesis–that driving resilience- or susceptibility-associated behaviors will change resilience–we used closed-loop DAT::NAc activation to reinforce fighting back or escape from attack. These behaviors were selected as resilient-associated and susceptible-associated on the basis of increased behavioral expression (fight-back, Fig. 1i) and on increased dopaminergic activation (fight-back and escape, escape defined as the offset of proximity following fleeing from attack; Fig. 3f,4c; see Extended Data Fig. 9b-d for stimulation details).

Fight-back triggered DAT::NAc activation increased fighting back behavior between early versus late defeat (Extended Data Fig. 9g, N=16, t=2.7, p=0.04, day 9-10 vs 1-2). Compared to controls, animals receiving fight-back triggered activation more often occupied the “close vigilance” cluster (5) behavior that we observed correlated with resilience in females (Fig. 5l, t=2.11, p=0.044) and had increased social interaction during the sensory period of defeat and in the SI test (Fig. 5m-n), indicative of the stimulation increasing aggressor value. Activation timed to fighting back also resulted in increased exploration (Fig. 5o-p) and mobility (Fig. 5q) relative to controls, suggesting increased resilience beyond social contexts.

Activation of DAT::NAc timed to a susceptibility-associated behavior (escape) significantly increased the likelihood of fleeing from attack from early to late defeat (Extended Data Fig. 9h, N=8, t=6.4, p<0.001, day 9-10 vs 1-2) and did not increase “close vigilance” behavior relative to controls (Fig. 5s). However, this reinforcement of escape behavior was not sufficient to increase susceptibility (Fig. 5t-x), suggesting either (1) that fleeing and its dopaminergic correlates are reflections of rather than causal drivers of susceptibility or (2) that aggressor value was increased by DAT::NAc activation within the defeat context, offsetting any pro-susceptible effects of DA reinforcement of avoidance behavior.

We next tested the hypothesis that decreases in DAT activity during defeat could lead to susceptibility by performing attack-triggered closed-loop inhibition of DA neurons in the VTA during defeat. Halorhodopsin (NpHR) was injected bilaterally in DAT::Cre animals and stimulation fibers were implanted over the VTA (Extended Data Fig. 9i). Expression was confirmed via real-time place aversion (Extended Data Fig. 9j). Surprisingly, closed-loop inhibition during defeat led to less time being attacked relative to controls (Extended Data Fig. 9o t=-3.76, p=0.001), as attack-inhibited animals were less likely to flee and thus promote attack (Extended Data Fig. 9p, t=-3.92, p<0.001). Attack-triggered inhibition did not lead to susceptibility in our post-hoc behavior tests (Extended Data Fig. 9q-u). Thus, while increasing DAT activity may bias towards resilience, decreased activity was not sufficient for susceptibility, potentially due to alterations in the defeat behavior. These data suggest that susceptibility may require both an increase in avoidance and a decrease in DAT activity during attacks.

The fact that both open loop and fight back triggered DAT::NAc stimulation produced resilience while similarly reorganizing defeat behavior toward more resilience-associated patterns (Fig. 5e,i,y) implies a common mechanism that is not critically dependent on the precise timing of DA with respect to actions. This common mechanism may be to increase the aggressor value (Fig. 5a,z), supported by several lines of behavioral and neural evidence. First, we find that NAc(DAT::GCaMP6f) correlates are consistent with a higher subjective value of the aggressor in resilient mice of both sexes (Fig. 3). Second, stimulation increased socially appetitive behaviors during and after defeat (i.e. more often occupying the “close vigilance” behavior cluster, higher SI time, closer average distance during sensory period, Fig. 5e-g, i-n). Lastly, this hypothesis is consistent with the well-established role of NAc DA in increasing stimulus values outside the social context^41,42,52,53^. Here we propose that such increased valuation may not only result in behavioral change towards the aggressor during and after defeat, but also produce a generalized resilience phenotype that extends beyond social approach to decreased anxiety-like behavior and increased exploratory behavior and mobility (Fig. 5h-i,o-q).

These results may help unite long-standing, divergent lines of evidence about whether increased DAT neuron activity is pro-resilient or pro-susceptible, by suggesting that the context of the manipulation determines its effects. While activating DAT neurons after chronic mild stress and *during* the stressor of tail suspension increased resilience (i.e. amount of struggling)^7^, previous work that activated DAT::NAc neural activity *after* stressful social interactions (when the barrier was replaced on defeat days or during the SI test the day after defeat) reported increased susceptibility rather than resilience.^4^ Post-defeat stimulation may have biased towards susceptibility by mimicking DA correlates of relief following aggressor interactions (Fig. 3d-f), but on a longer timescale. Consistent with this interpretation, we observed that the relative quantity of NAc(DAT::GCaMP6f) activity in the period directly after versus during defeat encounters was higher in susceptible than resilient mice (Extended Data Fig. 10). In addition, susceptible individuals have been shown to have enhanced spontaneous firing rate of DAT neurons following defeat.^3,5^ However, we cannot rule out that some details in our paradigm (e.g. behavior tested in the active rather than resting period of the light cycle) may have contributed to the pro-resilient rather than pro-susceptible effect of DAT neural stimulation.

In total, our work highlights the power of combining complex, ethologically relevant social stress paradigms with tools for behavior quantification^54^, chronic and projection-specific neural recording, and behavior-specific neural manipulations to improve our understanding of the role of neuromodulation in mediating individual differences in pathological outcomes.

---

**Extended Data Fig. 1.**
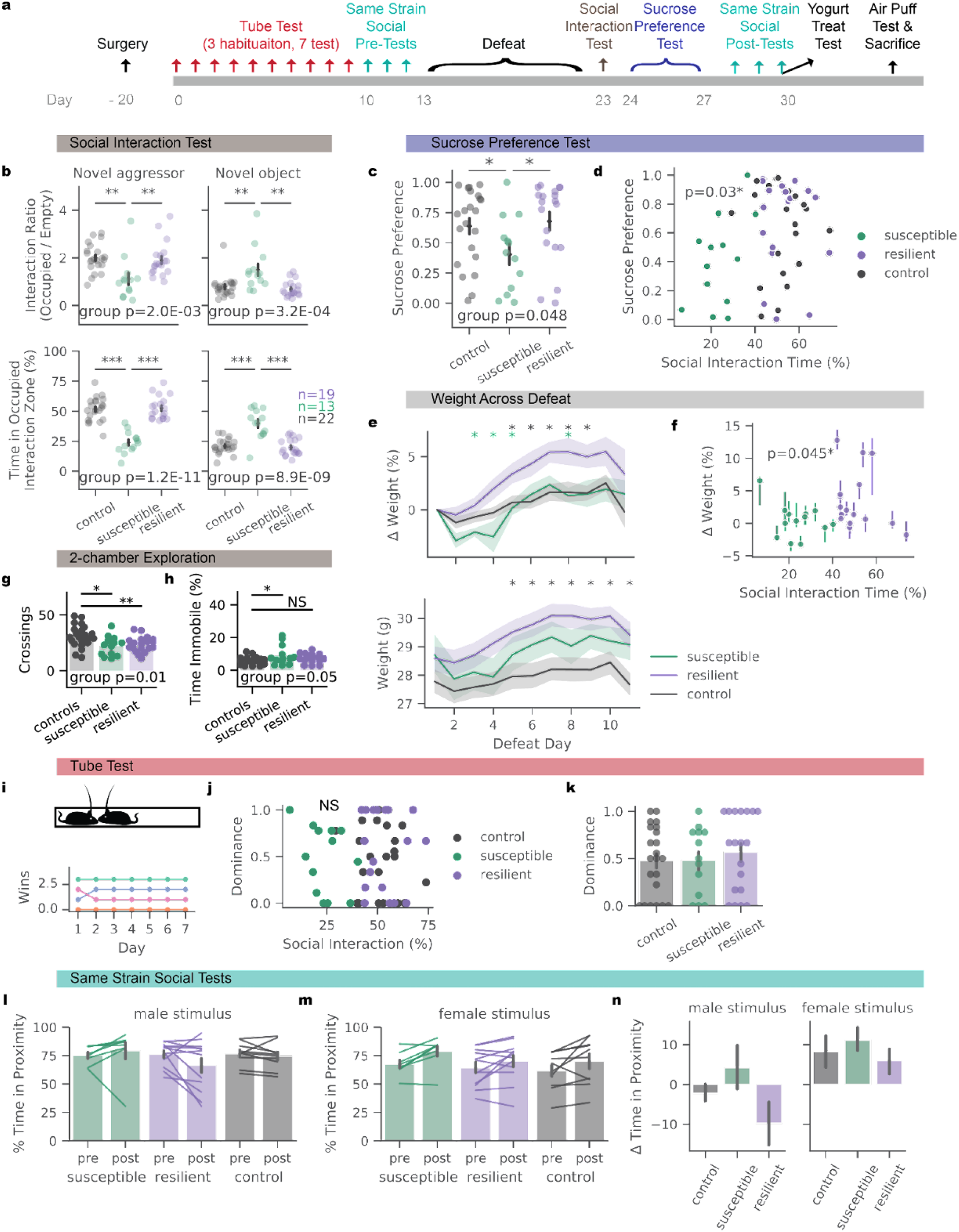
Pretests do not predict susceptibility while additional post-hoc tests confirm resilient and susceptible phenotypes. **a**, Timeline of behaviors for mice undergoing behavioral and neural recordings across chronic social defeat stress. **b**, In the social interaction test, social interaction ratio and time (N=22 controls, 13 susceptible, 19 resilient in **b**-**k**; 1-way ANOVA for social interaction ratio with mouse category as factors, F_(51,2)_=7.015, p=0.002, t-Test control vs susceptible p=0.001, resilient vs susceptible p=0.009, control vs resilient p=0.646; 1-way ANOVA for social interaction social interaction time with mouse category as factors, F_(51,2)_=42.9, p=1.18E-11, t-Test control vs susceptible p<0.001, resilient vs susceptible p <0.001, control vs resilient p=0.9). In social interaction test, object interaction ratio and time (1-way ANOVA object interaction ratio with mouse category as factors, F_(51,2)_=9.442, p<0.001; group effects of control vs susceptible, t=3.71, p=0.001; t-Test control vs susceptible p=0.003, resilient vs susceptible p=0.001, control vs resilient p=0.5; 1-way ANOVA for object interaction time with mouse category as factors, F_(51,2)_=27.26, p=8.86E-9; t-test control vs susceptible p<0.001, resilient vs susceptible p<0.001, control vs resilient p=0.6). **c**, Fraction of sucrose over total volume consumed in a 2-bottle sucrose preference test (1-way ANOVA for sucrose preference with mouse category as factors, F_(51,2)_=3.21, p=0.048; t-Test control vs susceptible p=0.039, resilient vs susceptible p=0.024, control vs resilient p=0.9). **d**, Relationship between sucrose preference and SI time (Pearson correlation coefficient R=0.29, p=0.03). **e**, Change in weight across defeat (top), absolute weight (bottom) (t-test for difference of means between resilient vs. susceptible and resilient vs. control groups, FDR-corrected for 20 comparisons in each sub-panel, threshold for significance of p<0.05, green stars indicate resilient vs susceptible and black indicate resilient vs control significance). **f**, Relationship between SI time and the change in weight during defeat (GEE, grouped by mouse, z=2.01, p=0.045). **g,** Number of crossings between chambers of a 2-chamber arena (1-way ANOVA for crossings with stimulation groups as factors, F_(51,2)_=5.77 p=0.006; post-hoc t-Test susceptible vs control p=0.011 and resilient vs control p=0.008). **h**, Percent of time immobile (speed < 1cm/s) during exploration of a 2-chamber arena (1-way ANOVA for crossings with stimulation groups as factors, F_(51,2)_=3.16 p=0.05; post-hoc t-Test susceptible vs control p=0.032 and resilient vs control p=0.071). **i**, Schematic of tube test for social hierarchy (top), example of tube test wins for a homecage of mice (colored by individual) measured across 7 days. **j**, No relationship between dominance (percentage of wins in the last 3 days of the tube test) and social interaction time in the social interaction test (N=22 controls, 13 susceptible, 19 resilient; Pearson’s correlation coefficient R=-0.038, p=0.78). **k**, Summary of **j** (1-way ANOVA for social interaction social interaction time with mouse category as factors, F_(51,2)_=0.342, p=0.712). **l**, Time spent in proximity with a male of the same strain (C57/BL6), in a freely moving environment (2-way mixed ANOVA with resilience group and time point as factors: group x time F_(64,2)_=0.319, p=0.728) **m**, Time spent in proximity with a female of the same strain (C57), in a freely moving environment. (2-way mixed ANOVA with resilience group and time point as factors: group x time F_(64_,_2)_=0.189, p=0.828) **n,** Difference in time spent with a male or female social target after vs before defeat. N=11 control, 8 susceptible, and 16 resilient mice in **l**-**n**.

**Extended Data Fig. 2.**
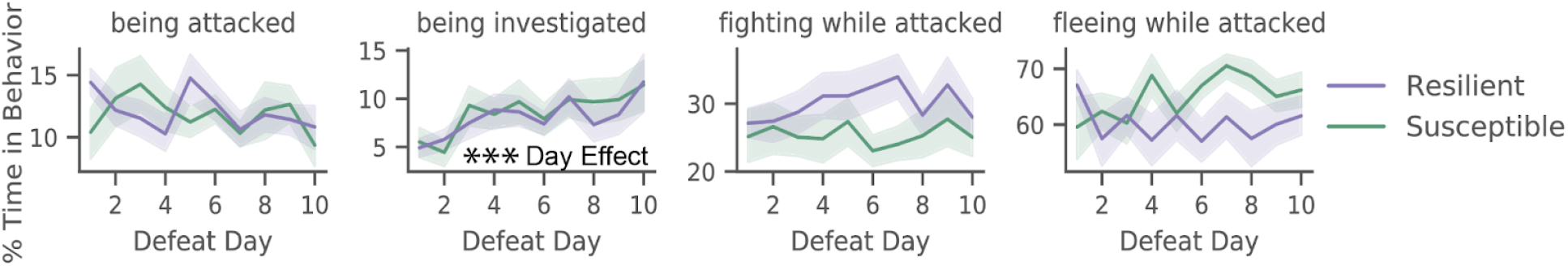
Evolution of defeat behaviors across defeat days. Amount of time spent in classified behaviors across days (GEE for time attacked: effect of resilience group z=0.27, p=0.79, effect of day z=-0.613, p=0.54; GEE for time investigated z=-0.031, p=0.98, effect of day z=4.01, p<0.001; GEE for fighting while attacked: effect of resilience group z=0.90, p=0.37, effect of day z=0.02, p=0.98, GEE for fleeing while attacked: effect of resilience group z=0.21, p=0.84, effect of day t=1.76, p=0.078, N=32 mice total, 19 resilient and 13 susceptible).

**Extended Data Fig. 3.**
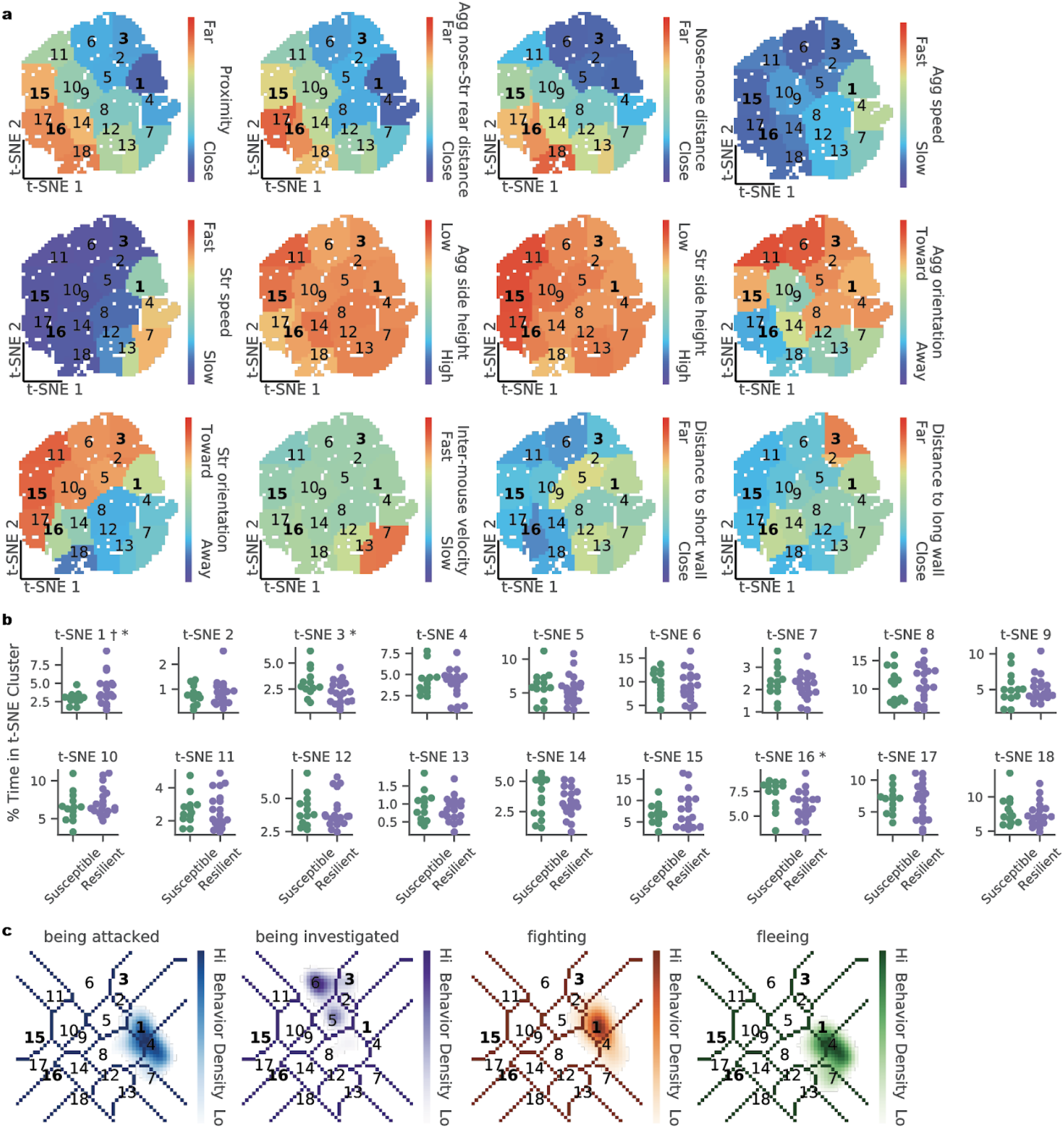
t-SNE clusters of social postures represent discrete behaviors, some of which are preferentially occupied by resilient or susceptible mice. **a**, Average raw feature value within each t-SNE cluster (scale in the full range of each variable, Str: Stressed mouse, Agg: Aggressor mouse). **b**, Average t-SNE cluster occupancies for each individual, split by susceptible and resilient groups (2-sample T-test *p<0.05; 2-sample Levene Test ^+^p<0.05, N=13 susceptible mice and 19 resilient). **c**, Density of behaviors annotated with supervised classifiers within t-SNE space, bold font indicating clusters that differentiate resilient and susceptible mice (Fig. 1o-r).

**Extended Data Fig. 4.**
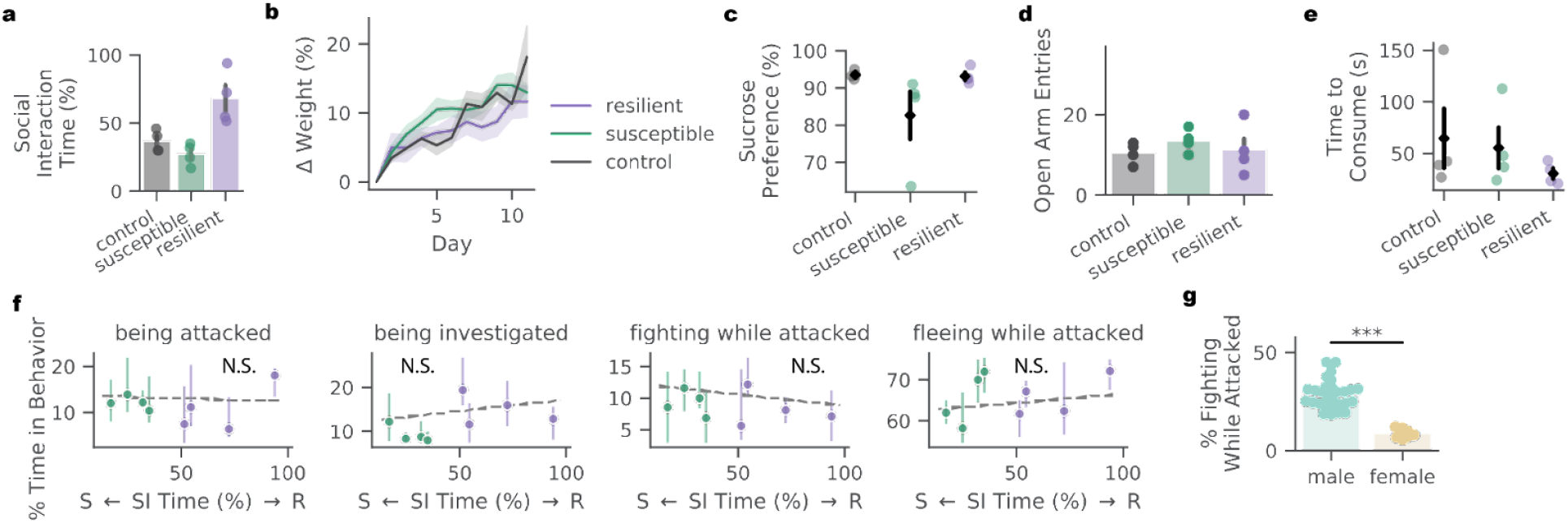
Behavioral and physiological differences across defeated females. **a**, Distribution of social interaction (SI) times in the post-hoc SI test (see Fig. 1b) in unstressed control and stressed females split into resilient and susceptible groups (N=4 controls, 4 susceptible, 4 resilient). **b**, Change in weight across defeat by females in control (N=4), resilient (N=4), or susceptible (N=4) groups. **c**, Fraction of sucrose over total volume consumed in a 2-bottle sucrose preference test. **d**, Number of entries into the open arms of an elevated plus maze. **e**, Time to consume a yogurt treat in novelty suppressed feeding assay. **f**, Time spent by individual mice in each behavior of interest (GEE: being attacked p=0.76; being investigated p=0.26; fighting while attacked, p=0.16; fleeing while attacked p=0.20; means plotted with 30th to 70th percentile error bands across the 10 defeat days; N.S., not significant). **g**, Difference in time males and females spent fighting back while attacked during defeat (2-sample t-Test, t=-7.45, p<0.001 N=32 males, 8 females).

**Extended Data Fig. 5.**
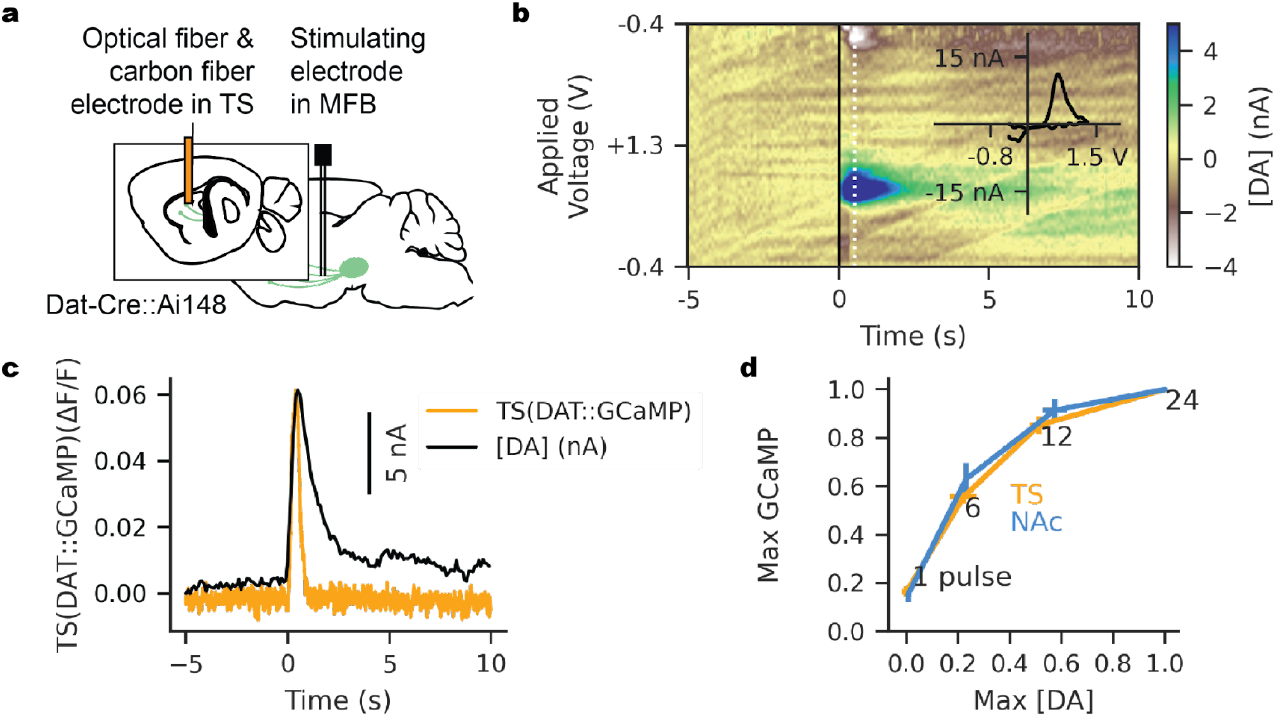
Simultaneous fast scan cyclic voltammetry and fiber photometry calcium recordings in the TS. **a**, Schematic for recording dopamine release and calcium fluorescence in the tail of the striatum (TS). **b**, Example 3-D color plot from a single recording site, where the medial forebrain bundle (MFB) was stimulated with 24 pulses at 60 Hz at time 0 (average of 5 trials shown). Inset shows the voltammogram at 0.5s after stimulation (white dotted line in color plot), which shows oxidation and reduction peaks consistent with the profile of dopamine. **c**, Time course of simultaneously measured calcium fluorescence and dopamine current (average over the same 5 trials shown in b). **d**, Calcium fluorescence and dopamine concentration peaks increase together with the number of pulses of stimulation delivered to the MFB (bars show standard error, N=6 recording sites). VTA-NAc data obtained with permission from Parker et al 2016 (N=4 recording sites).

**Extended Data Fig. 6.**
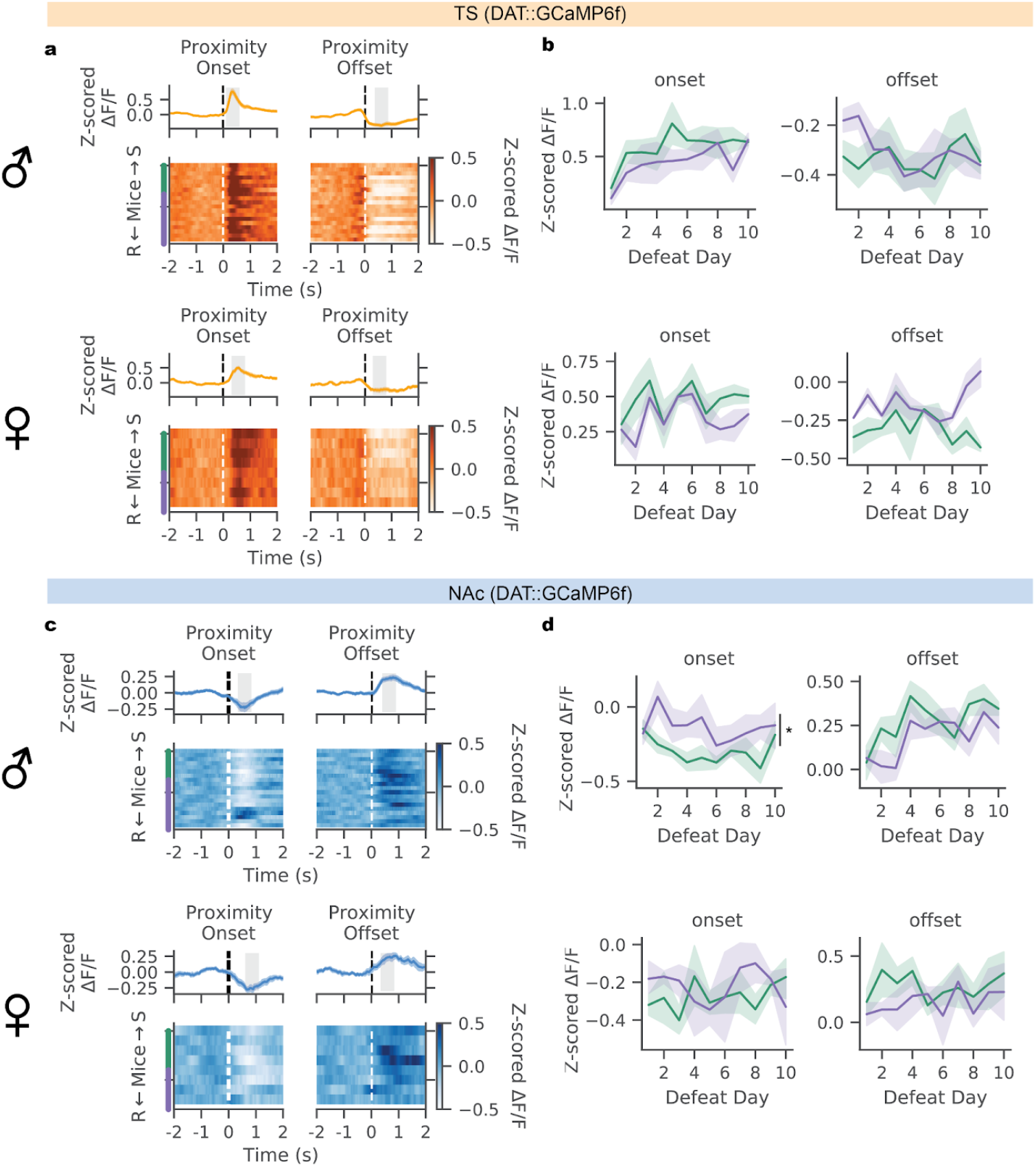
DAT::GCaMP6f activity profiles during defeat are similar in males and females. **a**, Average proximity onset- and offset-aligned TS(DAT::GCaMP6f) activity across stressed males (top) and females (bottom). **b**, TS(DAT::GCaMP6f) activity in the 0.5s at the onset or offset of social encounters (gray shaded zones in **a**) in males (top) and females (bottom) from resilient and susceptible groups across the 10 days of defeat. Solid line shows mean across mice and shaded region shows standard error. (GEE regression for male onset activity: main effect of day Z=5.71, p<0.001; resilience Z=-0.96, p=0.337; GEE regression for male offset activity: main effect of day Z=0.45, p=0.65; resilience Z=0.924, p=0.355; GEE regression for female onset activity: main effect of day Z=0.673, p=0.501; resilience Z=-0.955, p=0.34; GEE regression for female offset activity: main effect of day Z=-1.1, p=0.3; resilience Z=0.64, p=0.52; interaction between day and resilience Z=2.326, p=0.02). **c**, Same as **a** for NAc(DAT::GCaMP6f). **d**, Same as **b** for NAc(DAT::GCaMP6f) (GEE regression for male onset activity: main effect of day Z=-1.328, p=0.184; resilience Z=2.446, p=0.014; GEE regression for male offset activity: main effect of day Z=4.966, p<0.001; resilience Z=-1.34, p=0.18; GEE regression for female onset activity: main effect of day Z=1.369, p=0.171; resilience Z=1.578, p=0.114; GEE regression for female offset activity: main effect of day Z=0.041, p=0.986; resilience Z=-1.621, p=0.105).

**Extended Data Fig. 7.**
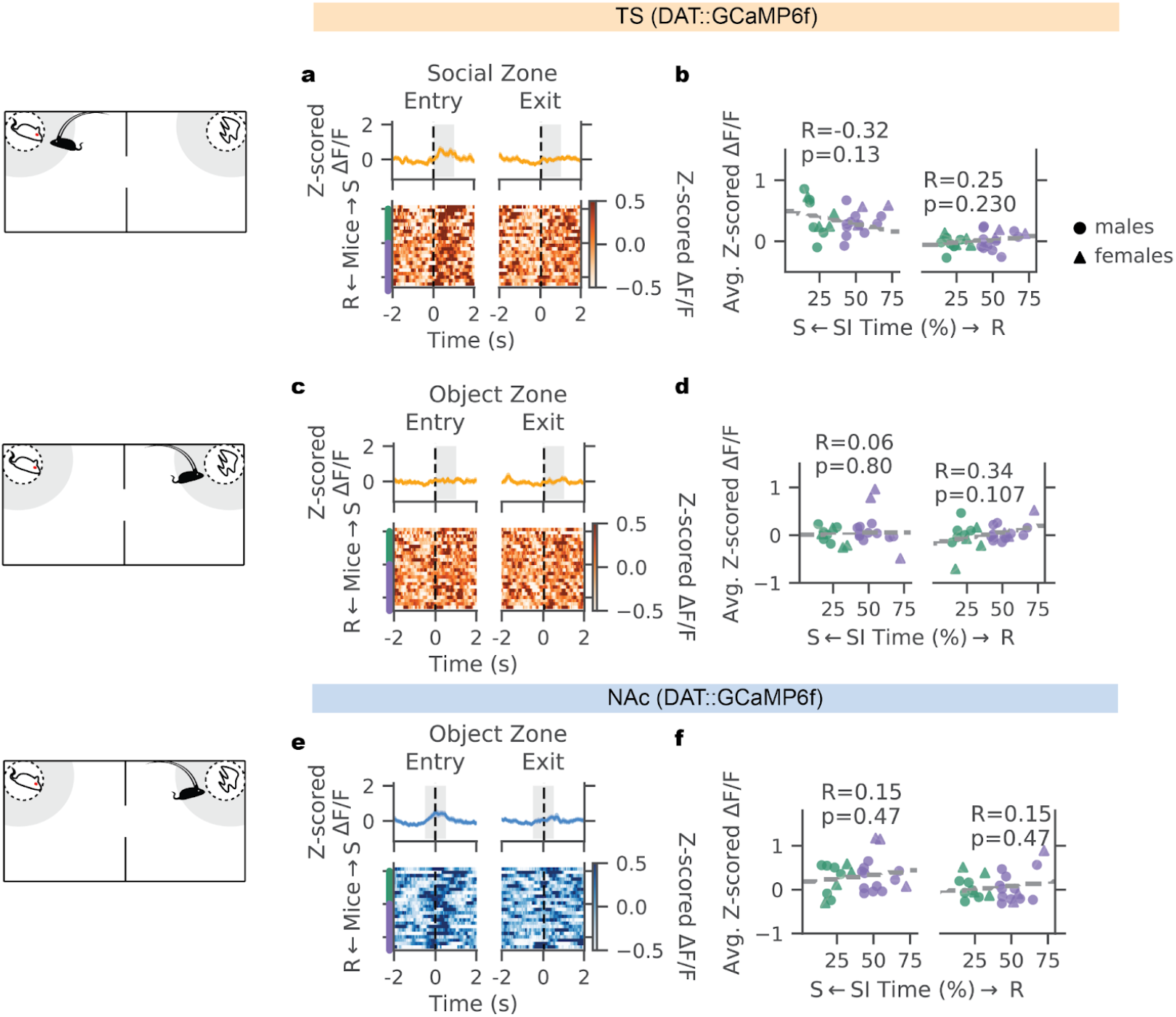
During the SI test, DAT::GCaMP6f activity in TS during social proximity and activity in TS and NAc in object proximity are not associated with resilience. **a**, Average social interaction zone entry- and exit-aligned TS(DAT::GCaMP6f) activity across stressed animals (mean plotted with standard error, N=6 susceptible males, 4 susceptible females, 10 resilient males, 3 resilient females). **b**, Relationship between individuals’ SI time and average TS(DAT::GCaMP6f) activity in the 1s after entry into the social interaction zone (left, Pearson correlation, R=-0.32 p=0.13) and in the 1s after exit from the social interaction zone (right, Pearson correlation, R=0.25, p=0.23). **c**, Average object zone entry- and exit-aligned TS(DAT::GCaMP6f) activity across stressed animals. **d**, Relationship between individuals’ SI test interaction time and average TS(DAT::GCaMP6f) activity in the 1s after entry into and exit from the object interaction zone (entry: Pearson correlation, R=0.06 p=0.8; exit: Pearson correlation, R=0.34, p=0.107). **e**, Same as in **c** for NAc(DAT::GCaMP6f). **f**, Relationship between individuals’ SI test interaction time and average NAc(DAT::GCaMP6f) activity in the 1s surrounding entry into and exit from the object interaction zone (entry: Pearson correlation, R=0.15 p=0.47; exit: Pearson correlation, R=0.15, p=0.47).

**Extended Data Fig. 8.**
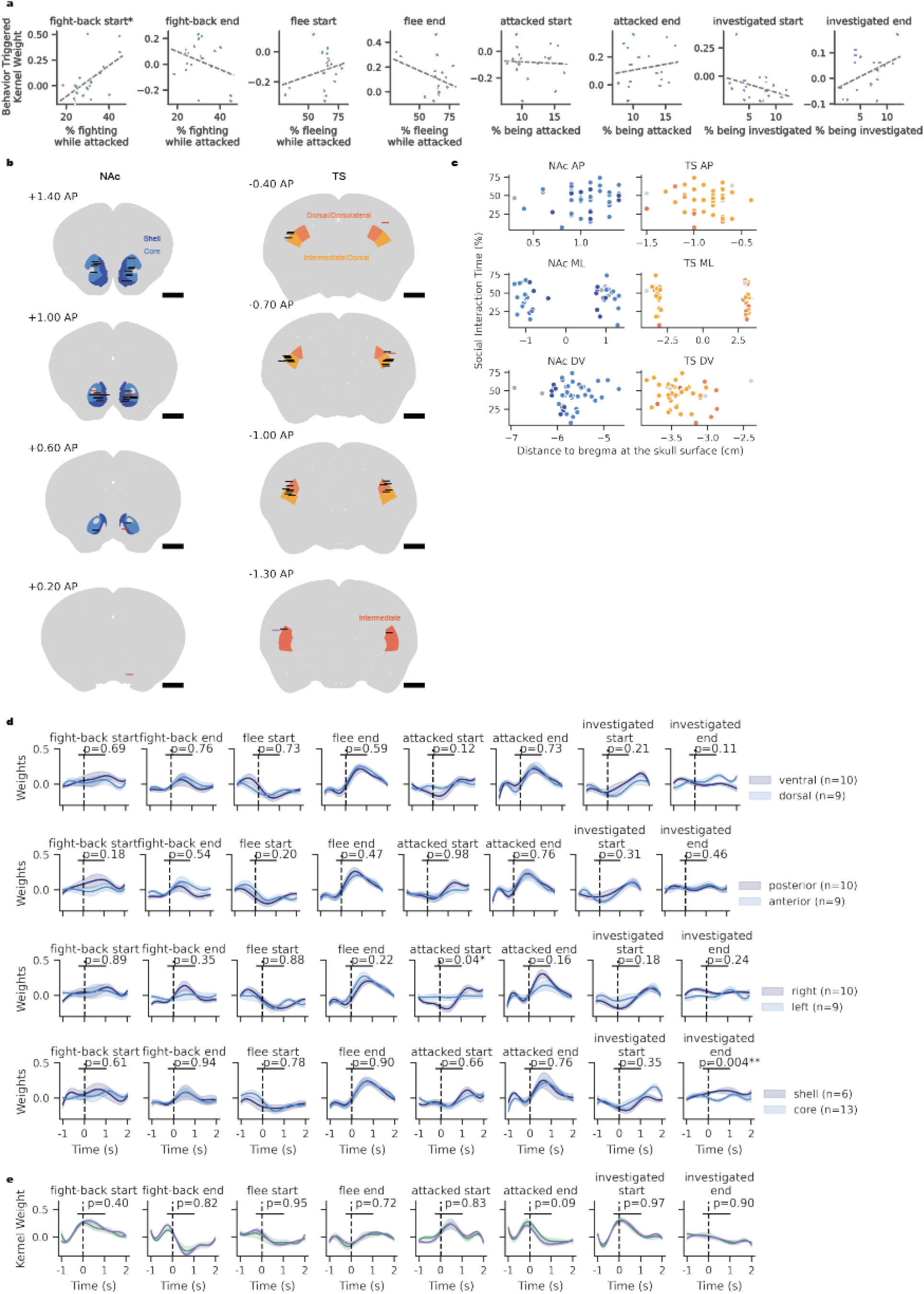
Fiber location for photometry recording and behavior encoding by the TS does not predict resilience in males. **a**, Relationship between average time spent engaging in each behavior and kernel weight for NAc(DAT::GCaMP6f) activity encoded by corresponding behavioral events (Pearson correlation, *p<0.05, N=19 mice, purple designating resilient animals, green showing susceptible). **b**, Anatomical locations of photometry fiber tips targeted to the nucleus accumbens (NAc) and tail of the striatum (TS), overlaid on regions defined by the Paxinos Atlas (scale bar 1mm, Paxinos splits TS into subregions of the caudal caudoputamen). Animals omitted on the basis of poor signal shown in red, or omitted on the basis of poor targeting shown in magenta, included animals shown in black. **c**, Lack of relationship between location of recording fiber tips and SI time. **d**, Average kernel weights for behaviors encoding NAc(DAT::GCaMP6f) activity in mice grouped by fiber location in the NAc (p-values shown for t-test for difference in means in average activity from 0.5 to 1.5s following the behavior event). **e**, Average kernel weights for behaviors encoding TS(DAT::GCaMP6f) activity in mice grouped by susceptibility (p-values shown for t-test for difference in means in average activity from 0.5 to 1.5s following the behavior event, N=7 susceptible (green) and 12 resilient (purple) mice).

**Extended Data Fig. 9.**
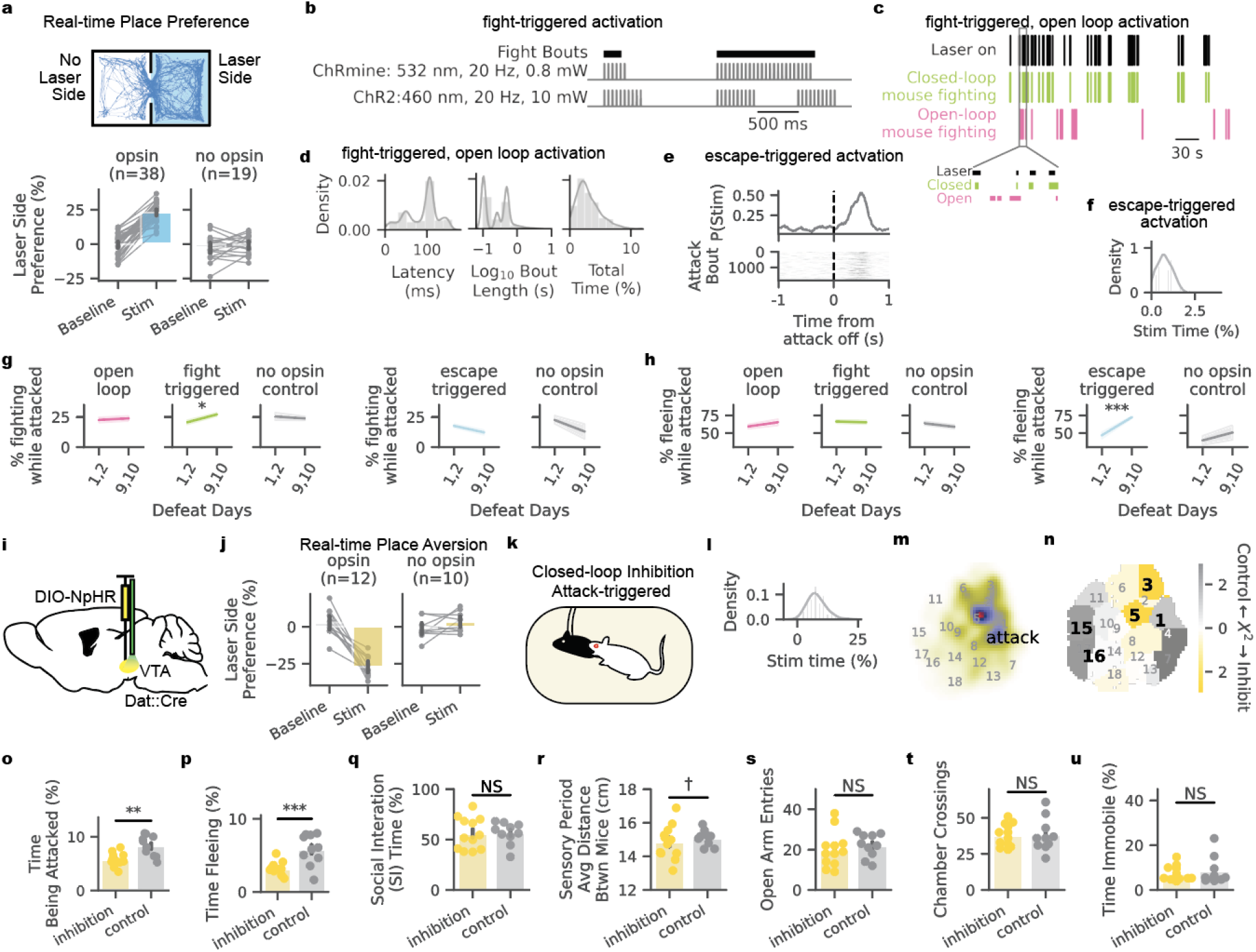
Additional characterization of optogenetic manipulation of DAT activity during defeat. **a**, Arena schematic and example stimulation session trace from a 2-chamber real-time place preference assay. 20-Hz laser stimulation was delivered whenever a mouse’s centroid was in the “laser side” of the chamber, sides counterbalanced across mice (top). Preference for the laser-stimulation side of a chamber in mice expressing or not expressing excitatory opsin. No stimulation was delivered in the baseline condition (bottom). **b**, Animals receiving fight-back-triggered optogenetic activation expressed either ChRmine or ChR2 and were delivered 20 Hz laser stimulation with the following parameters: ChRmine mice received green laser stimulation in sets of 3 pulses without pause for the duration of fighting bouts while ChR2 mice received blue laser stimulation in sets of 10 pulses with a 0.5s pause between pulse sets for the duration of fighting bouts. **c**, Example of a defeat session with laser light delivery triggered on fighting of the closed-loop mouse and simultaneously delivered to his open-loop neighbor, bottom inset is a 10s segment. **d**, Statistics across fight-back-triggered stimulation: Distribution of latency to trigger stimulation from capture of a fighting video frame, with a median of 100ms (left). Distribution of durations of stimulation bouts, 94% of which were less than 1s (middle). Distribution of the amount of defeat time in which light was delivered (mean is 2.99% of the defeat session) (right). **e**, Attack-offset aligned laser stimulation during escape-triggered activation. **f**, In escape-triggered activation, distribution of the amount of defeat time in which light was delivered (mean is 0.75% of the defeat session). **g**, Time spent in random forest labeled fighting behavior early (days 1,2) versus late (days 9,10) in defeat. (Paired t-Tests: 3-way FDR corrected: open-loop t=0.38 p=0.71, fight-triggered t=2.77 p=0.04, no opsin t=-0.41 p=0.71; 2-way FDR corrected: escape-triggered t=-2.33, p=0.10, no opsin t=-1.14, p=0.34). **h,** Average expression of random forest labeled fleeing behavior early (days 1,2) versus late (days 9,10) in defeat. (Paired t-Tests: 3-way FDR corrected: open-loop t=1.12 p=0.42, fight-triggered t=-0.29 p=0.78, no opsin t=-1.49 p=0.42; 2-way FDR corrected: escape-triggered t=6.42, p<0.001, no opsin t=0.68, p=0.54). **i**, Schematic of inhibition of dopaminergic cell bodies in the VTA (Dio-NpHR injection and fiber implant over the VTA of DAT::Cre animals). **j**, Real-time place aversion assay, with constant 7 mW green light delivery on the stimulation side of the chamber. Preference for the non-stimulation side of a chamber in mice expressing or not expressing inhibitory opsin. No stimulation was delivered in the baseline condition. **k**, Schematic for closed-loop, attack-triggered inhibition during defeat. **l**, In attack-triggered inhibition, distribution of the amount of defeat time in which light was delivered (mean is 7.93% of the defeat session). **m**, Density map of where in t-SNE behavior space animals were when receiving inhibition. **n**, Difference in occupancy of t-SNE clusters between inhibition (N=12) and no opsin control (N=10) groups, measured by individual clusters’ chi^2^ statistics; black bold numbers label behaviors differentially expressed by resilient and susceptible mice in our observational study. **o**, Average time across 10 days of defeat being attacked (t-Test, attack-triggered inhibition vs no opsin control, t=-3.03, p=0.0066). **p**, Average time across 10 days of defeat fleeing (t=-3.92, p=0.0008). **q**, Social interaction (SI) time in the SI test (t-Test, difference in SI time between inhibition and no opsin control groups: t-Test, t=0.04, p=0.96). **r**, Average distance between stressed and aggressor mice during the first 5 minutes of the barrier-separated sensory period across the 10 days of defeat (Levene test for difference in variance, W=4.92, p=0.038). **s**, Number of entries into the open arms of an elevated plus maze (t-Test, difference in mean entries between inhibition and no opsin control, t=-0.64, p=0.53). **t**, Number of crossings between chambers of 2-chamber arena (t-Test, difference in mean crossings between inhibition and no opsin control groups: t=-0.024, p=0.98). **u**, Percent of time immobile (speed < 1cm/s) during exploration of a 2-chamber arena (t-Test, difference in mean immobility between inhibition and no opsin control groups: t=-0.47, p=0.64).

**Extended Data Fig. 10.**
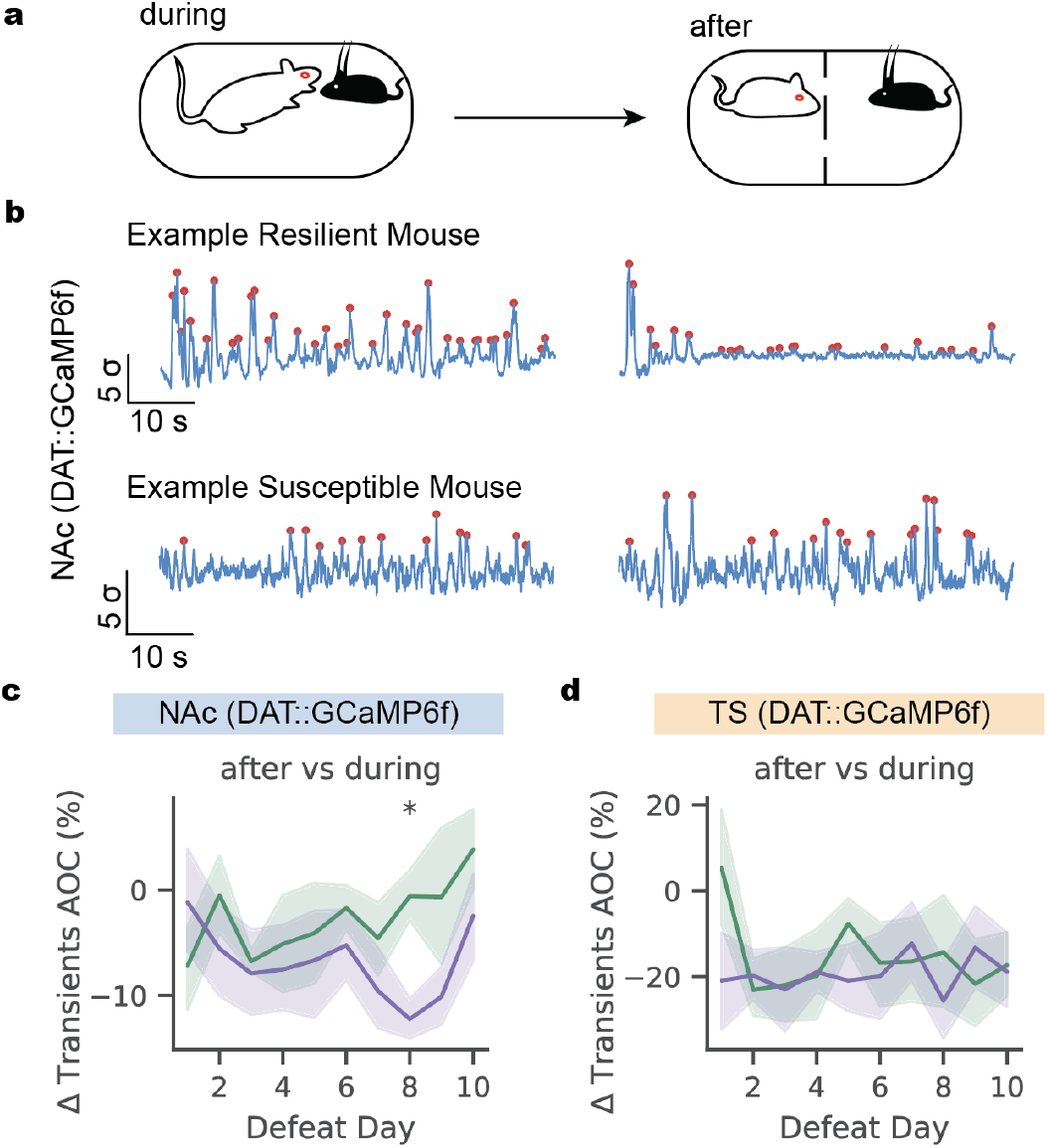
Difference in NAc(DAT::GCaMP6f) activity after versus during direct contact phase of defeat is higher in susceptible compared to resilient individuals. **a**, Schematic showing phases of defeat stress during and after direct contact. **b**, Example traces of NAc(DAT::GCaMP6f) activity during (left) and after (right) the direct contact phase of defeat from day 10 in an example resilient mouse (top) and susceptible mouse (bottom). Transients represented by red dots. Area under the curve (AOC) in next panels refers to the integral between the half max surrounding each transient. **c**, Difference in NAc(DAT::GCaMP6f) after vs during the contact phase of defeat (Mixed linear model for difference in AOC by day, resilience category, and their interaction; interaction effect z=-2.037, p=0.042; t-Test for difference in ratio of activity in susceptible vs resilient individuals, day 8: t=-3.607, p=0.02 after Bonferroni correction for 10 tests across 10 days). **d**, Difference in TS(DAT::GCaMP6f) after vs during the direct contact phase of defeat (Mixed linear model for difference in AOC by day, resilience category, and their interaction; no significant effects).

## Methods

### Mice

All animal procedures were approved by the Princeton University Institutional Animal Care and Use Committee and were in accordance with National Institutes of Health standards.

One hundred fifty experimental animals were used in this study. In behavior and fiber photometry recording studies, we used 10 wild type C57BL/6J males, 40 Dat::Cre (Jackson Laboratory strain 006660) crossed with the Ai148 GCaMP6f reporter line Jackson Laboratory strain 030328^55,56^) males, 6 Dat::Cre males, and 12 Dat::Cre x Ai148 females. In fast scan cyclic voltammetry, we used 3 Dat::Cre x Ai148 mice. For optogenetics, we used 70 Dat::Cre males and 9 wild type males. The above mice were naive to non-littermates.

Male and female conspecifics used as social targets in freely moving assays were sexually naive, C57BL/6J mice between the ages of 8 and 26 weeks. Aggressor animals used to deliver chronic social defeat stress were WT Swiss Webster or Esr1::Cre backcrossed to Swiss Webster strain breeders or retired breeders between the ages of 8 and 57 weeks. Mice were housed in a 12 h light-dark cycle with experiments taking place exclusively during the dark phase. Food and water were given *ad libitum*. Prior to behavioral testing, mice were group-housed with 2-5 mice per cage. All animals were bred within the Princeton University animal facilities or purchased from Taconic Biosciences.

### Surgery

At 6-12 weeks of age, animals were anesthetized (isoflurane at 3-5% for induction and 1-3% for maintenance) and leveled in a stereotaxic frame before proceeding with injections and implants specific to recording or manipulation experiments.

For GCaMP6f fiber photometry recordings, Dat::Cre x Ai148 mice were implanted with 400 μM optical fibers in the TS (AP −1 mm, ML +/− 3.25 mm, DV −2.5 mm relative to the skull surface at Bregma) and NAc (AP 1.2 mm, ML +/−1 mm, DV −4.5 mm relative to the skull surface at Bregma), one on each hemisphere with hemisphere’s randomly selected (MFC_400/430-0.48_3mm_MF2.5_FLT for TS, MFC_400/430-0.48_6mm_MF2.5_FLT for NAc from Doric Lenses Inc.). Fibers were fixed to the skull with Metabond (Parkell). Then, titanium headplates^57^ were secured to the animals’ skulls (Ortho-Jet Crystal mixed with carbon glassy, spherical powder, Sigma-Aldrich). Animals were allowed to recover for a minimum of 7 days.

For optogenetic activation, Dat::Cre animals were injected with 500nL of viral vector expressing an excitatory opsin (ChR2: AAV5-EF1a-DIO-hChR2(H134R)-EYFP/mScarlet-WPRE-HGHpA or ChRmine: AAV5-EF1a-DIO-ChRmine-EYFP/mScarlet-WPRE-HGHpA; produced by the PNI virus core; injected at titers of 7E13 or 5E13 genome copies/mL) in the ventral tegmental area (VTA, AP −3.1 mm, ML +/−0.5 mm, DV −4.71 mm form the skull surface at bregma). In the same surgery or at least 4 weeks later, animals were implanted bilaterally with 200 μM diameter flat optical fibers or 100 μM diameter tapered optical fibers above or within the NAc respectively (AP 1.2, ML +/1.0, DV −3.9 to −4.2 mm from the skull surface at bregma, inserted at a 10° angle). No-opsin control animals of the same age (Dat::Cre or negative genotypes), were injected with a virus for EYFP (AAV5-EF1a-DIO-EYFP-WPRE-hGHpA, Addgene, titer: 2E13 genome copies/mL) or received no injection. Then, these animals received optical fiber implants as described for the opsin-expressing animals. Animals were allowed to express for a minimum of 8 weeks before behavioral experiments were performed.

For optogenetic inhibition, Dat::Cre animals were injected with 500nL of viral vector expressing the inhibitory opsin halorhodopsin (AAV5-EF1a-DIO-eNPHR3.0-EYFP-hGHpA; produced by the PNI virus core; inject at a titer of 4.4E13) in the ventral tegmental area (VTA, AP −3.1 mm, ML +/−0.5 mm, DV −4.71 mm form the skull surface at bregma). In the same surgery, animals were implanted bilaterally with 200 μM diameter flat optical fibers over the VTA (AP −3.1 mm, ML +/−1.6 mm, DV −4.2 mm form the skull surface at bregma, inserted at a 10° angle).

For induction of aggression toward females, Esr1::Cre Swiss Webster males were injected with a viral vector expressing excitatory DREADDs (Elf1a-DIO-hM3Dd(Gq)-mCherry, Addgene 50460; produced by the PNI virus core; titer of 1E14). Bilateral injections of 160 nl were targeted to the ventromedial hypothalamus ventrolateral area (VMHvl, AP −4.65 mm, ML +/−0.7 mm from the anterior sinus and DV −5.8 mm from the brain surface). Incisions were sutured and animals were allowed to recover and virus to express for a minimum of 3 weeks.

### Video and fiber photometry DAQ

For the tube test, chronic social defeat stress, freely moving object and social interaction tests, and free food reward assay (described below), fiber photometry and videography (BlackFly S camera, FLIR recorded with SpinView software) recordings took place with analogue signals monitored simultaneously through a RZ5P data acquisition board (Tucker Davis Technologies).

Specifically, from the camera, we recorded the analogue output signal indicating open exposure (via GPIO). The camera was triggered by the recording software at a rate of 120 frames per second (FPS) for chronic social defeat stress and 40 FPS for all others. The camera was mounted 2 inches above and 27 inches in front of the recording platform oriented at 90 degrees toward the side of the preparation. For assays other than the tube test, the camera also captured the top-down view of the preparation with a mirror mounted at a 40 degree angle above horizontal.

GCaMP6f recordings were made through a Doric Lenses phometry system (4-channel driver LEDD_r, LEDs at 465 and 405 nm, fluorescence mini cube FMC5_E1(465-480)_F1(500-540)_E2(555-570)_F2(580-680)_S, and Newport Visible Femtowatt Photoreceiver Module NPM_2151_FOA_FC). The system was driven by and recorded from using custom code written for a real-time processor (RZ5P, Tucker Davis Technologies). GCaMP was excited by driving a 465 nm LED light (~400 Hz at an intensity of ~10 μW, filtered between 465 and 480 nm) delivered to the brain through a fiber optic patch cord (MFP_400/430/1100-0.48_2m_FCM-MF2.5), with a separate LED and cable for each recording site (NAc and TS). The emission fluorescence passed from the brain through the same patch cords and was filtered (500-520 nm), amplified, detected, and demodulated in real time by the system. Demodulated fluorescence signals were saved at a rate of ~1 kHz.

Analog camera shutter signals were captured by the processor for synchronization of video and photometry signals.

Separate recordings were made during each behavior period: 1-10 minutes during the tube test, 5 minutes during chronic social defeat stress before the barrier was raised, 5-7 minutes during defeat while the barrier was raised and attack occurred, 5 minutes during defeat while barrier was returned, 5 minutes during same strain social interactions or object interaction, 3-10 minutes during free food reward, and 5-15 minutes during head-fixed air puff delivery.

### Behavioral assays

Animals undergoing fiber photometry recordings were subject to the following assays in this order: tube test, chronic social defeat stress, 2-chamber exploration, social interaction test, sucrose preference test, freely moving object and same strain social interaction test, free food reward, (novelty suppressed feeding in females), and head-fixed air puff. Animals undergoing optogenetic studies were subject to the following assays in order: real-time place preference or aversion (with at least 1 week delay until the next assay), freely moving object and same strain social interaction test, chronic social defeat stress, 2-chamber exploration, social interaction test, sucrose preference test, freely moving object and same strain social interaction test, and elevated plus maze test.

### Tube test

We used the tube test^58^ to establish social rank within the homecage. Prior to experiencing chronic social defeat stress, 56 mice from 18 cages of 2-5 mice each were evaluated for their social rank within their homecage using the tube test as described previously^59^. For three days, mice were habituated to the recording platform and 12-inch long clear plexiglass tube (1.15 inch inner diameter without a ceiling slit or 1.5 inch inner diameter with an elevated floor and ceiling slit for implant access). To habituate, each mouse made 10 passes through the tube each day (5 from either end).

To test mice for their social rank, for 7 days following habituation, every pair of mice in each cage was placed into the tube such that they met in the middle; every mouse faced each cagemate once per day. From each pair, the loser was defined as the mouse who stepped with a back paw out of the tube and onto the platform first. The other mouse who remained in the tube was defined as the winner. The average percentage of wins per day on the last 3 days of the tube test was used to determine the mouse’s dominance score.

### Chronic social defeat stress

Experimental mice were placed in the shoebox home cages (#5 Expanded Mouse Cage 8.75 x 12.125 x 6.395 inches, Thoren Caging Systems, Inc.) of singly housed male Swiss Webster (SW) aggressors. For recording behavior, food was removed from the cage and the typical stainless steel lids were replaced with a custom cut sheet of clear acrylic with a hole or slit for patch cables to run through. Shoebox cages were placed on a platform with an angled mirror and video recordings were made as described above.

Behavioral and neural recordings were made for 5 minutes before the barrier was removed, and the SW proceeded to defeat the experimental mouse for 5-7 minutes during which time recording continued. Finally, the barrier was replaced for an additional 5 minutes of recording. Stressed animals remained housed opposite the barrier from the aggressor until the following day. For each of the next 9 days, the stressed mice were rotated to be paired with and experience defeat from a novel SW. Following the 10 days of defeat, aggressors were removed and all mice were singly housed in the shoebox cages through the remaining stages of the experiment.

Unstressed control animals were pair-housed with a perforated barrier separating the two mice. They were handled and their cages rotated each day for 10 days.

### Social defeat stress in females

Defeat procedures and recordings were conducted as above with the following differences in aggressors and overnight housing conditions: Female stressed animals experienced attack from SW aggressors expressing hM3Dd(Gq) in the VMHvl. Twenty to 40 minutes prior to defeat interactions, aggressors were injected with Clozapine N-oxide (CNO) dihydrochloride (Hello Bio Inc, HB6149) diluted in sterile saline to provide a 0.1mL injection at a concentration of ~2 mg/kg. Following behavior and neural recordings (5 minutes with barrier, 5 minutes direct interaction, 5 minutes with barrier), stressed females were removed from aggressor homecages and housed together, 2 per cage, separated by a perforated barrier. Unstressed controls were also housed in barrier-separated pairs. Aggressors and females were housed overnight in separate rooms.

### Social interaction test

On the next day following 10 days of defeat stress (day 11), mice were evaluated for resilience by the social interaction test. First stressed mice were placed in a two-chamber arena (22 inches x 9.5 inches) with two empty mesh pencil cups in the far left and right corners for 5 minutes. Then the stressed mouse was removed from the chamber, and a novel aggressor male SW mouse was placed in beneath one cup while a novel object (eg. a helping hand clamp) was placed beneath the other. The stressed mouse was then returned to the recording chamber for an additional 5 minutes. Animal location was tracked via Ethovision (Noldus). In particular, we quantified the time spent within the social and object interaction zones (2.5 inches from enclosure). The time spent in the social interaction zone was used to delineate resilient and susceptible mice. The cutoff was defined as 1 standard deviation below the mean social interaction time over the unstressed control group.

### Freely moving object and same strain social interaction

Before and after defeat, video and fiber photometry recordings were taken as mice freely interacted with novel C57/BL6 social targets of both sexes as well as a novel object (e.g. tape roll, 50 mL Falcon tube). Recordings took place on the same platform described above for recording defeat. Female social targets were in estrus, evaluated via vaginal cytology^60^. Behavior occurred in the same shoebox cages and recording platform used for defeat. After at least 5 minutes of habituation to the recording cage, mice were presented with the novel mouse or object. Video (40 FPS) and fiber photometry recordings were taken for 5 minutes, beginning just before the introduction of the stimulus.

### Free food reward

A piece of a palatable treat (mini yogurt drops, Bio-serve F7577, cut with a razor plate into eighths) was dropped into the mouse’s home cage. Videos were hand-scored with BORIS^61^ event logging software for approaching (the single food-directed walking bout preceding sniffing and eating), sniffing (nose near the treat without chewing observed), and consuming the food (chewing observed).

### Head-fixed air puff

Mice were head-fixed while resting in a sliding plastic burrow.^62^ They habituated to the burrow for at least 3 minutes until they were able to maintain a motionless, neutral position in the burrow. Air puffs (~15 psi, 6 to 10 puffs per animal) were delivered to the right whisker pad at manually delivered intervals with at least 10s between puffs. Animals were required to be in a motionless, neutral position in the sliding burrow before puffs were delivered.

### Sucrose preference test

Preference for 1% sucrose (in water) compared to plain water was assessed with a two-bottle choice assay conducted across 3 days, with bottle locations randomly assigned to the left or right of the home cage. The preference was measured as the ratio of sucrose solution to total liquid (water and sucrose) consumed, measured by the difference in the weight of the bottles before and after the 3 days of free consumption.

### 2-chamber exploration

Mice were placed in a two-chamber arena (22 inches x 9.5 inches) for 5 minutes. Animal location and speed were continuously tracked via Ethovision (Noldus). From these tracks, we quantified the number of crossings made between the 2 chambers (defined by centroid passing over the chamber midline) and the amount of time spent immobile (speed < 1cm/s).

### Novelty suppressed feeding

Females were food-restricted for 18 hours prior to being placed in the corner of a brightly lit, novel chamber (19.5 x 19.5 inches) with a single yogurt treat placed in the center of the camber on a plastic platform. Latency to consume was scored as the time between entering the chamber and biting or licking the treat in the center. After the first consumption bout, animals were placed back in their home cages with ad libitum food access.

### Real time place preference/aversion

Prior to optogenetic experiments during defeat and at least 8 weeks following viral injections and at least 2 weeks following fiber implantation, mice were tested for opsin expression by preference for the stimulated half of a 2-chamber arena (22 inches x 9.5 inches). Green (532 nm, 0.8 mW for ChRmine or 4 mW for NpHR animals) or blue (447 or 460 nm, 7-12 mW for ChR2 animals) lasers were connected to a commutator (Doric Lenses, FRJ_1×2i_FC-2FC_0.22), which led to 200 μM diameter patch cords that were fastened to the mice’s implants via plastic sleeves surrounded by black electric tape. Lasers were triggered by a BNC cable connected to the output channel of a Pulse Pal (https://open-ephys.org/pulsepal), which was programmed to deliver 5 pulses of 5 ms in duration at 20 Hz every 1s for excitatory opsin animals and continuous light for inhibitory animals.

First, animals explored the chamber for 5 minutes without receiving light stimulation. Then during a 15 minute test period, animals received light stimulation when they were on the designated half of the chamber (alternating sides with animals run in a random order). Mouse centroids were tracked using Ethovision (Noldus) in real-time. When the mouse’s centroid was detected in the stimulated side of the chamber, the software sent a trigger (through a BNC cable connected to the Pulse Pal input port) to run the stimulation protocol as long as the mouse was in the stimulated side. Stimulation was halted when the animal was on the opposite side of the chamber.

### Elevated plus maze

Following defeat, animals from the optogenetic study were placed in the center of an elevated plus maze (2 enclosed arms and 2 open arms, arms 30 inches long and 2.5 inches wide) facing into the open arm away from the experimenter. The animal was free to explore the maze for 7 minutes, while its centroid location was tracked via Ethovision (Noldus). The number of entries into the open arms of the maze was used as a proxy for lack of anxiety.

## Behavioral Analysis

### Behavioral Annotation

For supervised classification of behaviors in defeat, ground truth was determined by hand annotations of videos scored with BORIS.^61^ The following behaviors were annotated: stressed mouse being attacked (rapid forward movement of the aggressor such as a lunge or chase directed at the stressed mouse), stressed mouse being investigated (stationary or slow-moving posture with the aggressor animal’s nose close to or in contact with the stressed mouse), stressed mouse fighting back (facing and actively engaged with the aggressor as opposed to leaning back or simply standing near the aggressor), and stressed mouse fleeing (rapidly running or jumping away from the aggressor). The number of annotated frames included 120,983 being attacked frames out of 896,178 across 25 videos, 51,170 being investigated frames out of 591,396 across 17 videos, 21,337 fight back out of 59,1396 across 17 videos, and 77,206 fleeing frames out of 896,178 across 25 videos. Annotations were carried out blind to the resilience or susceptibility of the behaving mice.

### Marklerless pose tracking

DeepLabCut software version 2.0.6 was used for tracking the positions of the stressed and aggressor mice during defeat. The training set included 1000 frames from 70 videos across 21 mice from 3 separate defeat cohorts (as well as 37 outlier frames). The following points were tracked:

- TopStressNose
- TopStressRightEar
- TopStressLeftEar
- TopStressFiberBase
- TopStressTTI
- TopStressTTip
- TopAggNose
- TopAggRightEar
- TopAggLeftEar
- TopAggTTI
- TopAggTTip
- BottomStressNose
- BottomStressRightEar
- BottomStressLeftEar
- BottomStressFiberBase
- BottomStressRightForePaw
- BottomStressLeftForePaw
- BottomStressTTI
- BottomStressTTip
- BottomAggNose
- BottomAggRightEar
- BottomAggLeftEar
- BottomAggTTI
- BottomAggTTip
- TopDividerRight
- TopDividerLeft
- BottomDividerTopRight
- BottomDividerTopLeft

(TTip: Tail tip, TTI: Tail-torso interface, Stress: stressed mouse, Agg: aggressor) The training was run for 1.03 million iterations with default parameters (training frames selected by kmeans clustering of each video session in the training set, trained on 95% of labeled frames, initialized with ResNet-50, batch size of 4).

### Feature definition

To define the stressed mouse’s posture with respect to his environment and aggressor, we converted pose data to the following behavioral features:

1. Between centroid distance: euclidean distance between the midpoint between each mouse’s tail-body interface and nose, defined by the top-down view.
2. Distance between aggressor nose and stressed mouse rear
3. Distance between aggressor nose and stressed mouse nose
4. Between centroid velocity: instantaneous (every 10 frames or 0.083s) change in between centroid distance, median smoothed with a window of 0.175s.
5. Aggressor speed: instantaneous distance between centroid position every 10 frames, smoothed as above
6. Stressed mouse speed: same as above
7. Orientation of aggressor with respect to stressed mouse
8. Orientation of stressed mouse with respect to the aggressor
9. Height of the aggressor: side view nose Y position
10. Height of stressed mouse: same as above
11. Distance of stressed mouse from the closest short wall of the cage: based on top-down view
12. Distance of the defeat mouse from the closest long wall of the cage

### Feature preprocessing

Features were truncated to fall within the 1st and 99th percentile of all recorded data for each feature (in order to remove extreme outliers), smoothed across time with a Gaussian Filter of 0.25 s, and rescaled from −1 to 1 (sklearn.preprocessing.MixMaxScaler) within each session to account for variability in mouse size and mildly varying camera angle or height. We chose to rescale features so that no single feature dominates due to higher magnitude while maintaining the original feature distributions and their covariances, properties that would not be maintained if each feature were normalized independently to unit variance for example.

### Supervised behavior classification

To identify behaviors across our entire video dataset, we trained supervised random forest classifiers using hand annotated data. Behaviors of interest during defeat included being attacked, being investigated, fighting back, and fleeing. These each were classified by a separate binary random forest classifier (Scikit-learn). The training and testing set consisted of 8 videos each. Ground truth was determined by hand annotation (BORIS) for frames in which the behavior was occurring (see above).

For each classified behavior, the feature matrix included the 12 features above for each video frame. The objective matrix was a binary indicator of if the behavior was hand-annotated in that frame. The training set was composed of all the frames in which the behavior was present and a randomly selected equal number of frames where the behavior was absent. The classifier was trained with a max depth of 2 and 100 estimators.

The probability threshold for detecting behaviors was set to the most permissive possible without exceeding a false positive rate of 3% on the training set. Evaluation was conducted by plotting the receiver operator curve on the held out testing set. The false positive and true positive rates of classification were: 1.6% and 68% for being attacked, 2.7% and 60% for being investigated, 2.3% and 56% for fighting back, and 2.6% and 53% for fleeing.

### Unsupervised behavior embedding and clustering

To characterize behaviors as stereotyped actions repeated throughout time, we followed previous work in using a low dimensional embedding of the original features and defining behaviors as high density clusters in that low dimensional embedding^63^. To achieve dense clusters we embedded our behavior features using t-distributed stochastic neighbor manifold (t-SNE), which preserves small pairwise distances and thereby retains clustering of nearby points.

Generating this manifold involved a technique known as importance sampling, which allowed us to create a final embedding including behaviors that might be rare or nuanced, and thus underrepresented in a uniform sampling over time. Importance sampling includes two rounds of t-SNE embedding. First, ~12 thousand frames of behavior were uniformly sampled in time across all videos (N=348) analyzed for Fig 1. Those features were embedded into a 2-component t-SNE manifold (sklearn.manifold.TSNE with perplexity=100). The embedded space was binned into a 50 x 50 histogram, smoothed with a 2D gaussian kernel of 2.5 standard deviations and parcelated into 18 clusters via watershed (skimage.morphology.watershed) over the smoothed histogram. As t-SNE is non-parametric, a separate model was necessary to learn the mapping from the original 12-dimensional feature space to this 2D t-SNE space. A multi-layer perceptron (sklearn.neural_network.MLPRegressor, hidden layer size of 400×200×50 units) was used for this purpose, allowing data from every video frame to be represented in 2D t-SNE space, and thus to fall into 1 of the 18 clusters defined in this space.

To generate the final t-SNE embedding, we sampled ~10 thousand frames from these 18 clusters, with additional sampling of frames where aggressive encounters took place. To do so, we characterized the overlap between random-forest classified attack frames and the clusters in t-SNE space. From the cluster most overlapping with attack, we sampled 5 random frames from every defeat video. From the 17 remaining non-attack clusters, we sampled 2 random frames from every defeat session. Thus, from 32 animals undergoing defeat for 10 days, we sampled (2 * 17) frames from non-attack clusters and 5 frames from the attack cluster on each day for each mouse for a total of 32*10*(17*2+5)=12,480 frames. From these sampled frames, we embedded the 12-dimensional behavior features into 2-component t-SNE space. The full set of video frames were then mapped into this final t-SNE manifold via another multi-layer perceptron. Then a 2D histogram of that perceptron-mapped 2D data was smoothed with a gaussian kernel of 1.5 standard deviations and divided into 18 clusters again via watershed. Gaussian kernels in both t-SNE steps were chosen as the roundest number to yield 10-20 clusters from watershed clustering.

### Resilience classification by unsupervised behavior occupancy

To test if the amount of time spent in the 18 t-SNE-based defeat behavior clusters was predictive of resilience or susceptibility in individual mice, we assessed leave-one-mouse-out cross-validated accuracy of making binary resilience or susceptibility predictions using logistic regression classifiers (sklearn.linear_model.LogisticRegression, max_iter=5000, class_weight=‘balanced’, penalty=‘none’). For each mouse, a single 1×18 vector was generated where each element represented the average fraction of time (across the 10 defeat days) in which that mouse occupied the corresponding t-SNE cluster. To avoid collinearity due to the 18 fractions summing to 1, only the first 17 clusters were used in the logistic regression training matrix. In each cross-validation round, the training matrix included an intercept and a chi-square-regularized set of 4 of the 17 t-SNE clusters (the 4 clusters with the highest univariate chi-square statistic separating resilient and susceptible mice in the training set, sklearn.feature_selection.chi2). For each held out mouse the training matrix was thus 5 (features) x 31 (mice). For each mouse, there was a new model for a total of 32 models, each with a different held out mouse. The accuracy was reported as the percent of accurately classified held out mice. The final model accuracy was 81.25% compared to 50%+/−8.5% (median and standard deviation) accuracy in “correctly” classifying data with shuffled resilience and susceptibility labels.

### Processing of calcium data

Each recording session was separately filtered and z-scored before further analysis. Demodulated fluorescence signals were processed by removing the linear trend (scipy.signal.detrend; removes downward linear trend due to fiber bleaching), bandpass filtering (0.1 to 30 Hz, scipy.signal.filtfilt with an order of 3; removes fluctuations outside of the timescale of GCaMP6f dynamics), normalizing by a moving average (window of 10 s; converts signal from fluorescence to change in fluorescence over a moving baseline, i.e. ΔF/F), and Z-scoring by the standard deviation of the detrended, filtered, normalized signal (normalizes the dynamic range across individuals to allow for comparison between individuals).

Poor quality recordings (n=47/259 sessions) were removed from further analysis by detecting the fraction of each session in which activity peaks occurred (Z-scored signal above 2) and defining poor quality recordings as those with activity peaks occurring below 2 standard deviations from the average amount of peak activity across sessions (from included animals). If the majority of an animal’s recording sessions were excluded, we excluded the remaining sessions from that animal (n=5/26 mice). Animals excluded from neural recording analysis were still included in behavioral analyses. Animals with poor targeting identified by registering extracted brains to the Paxinos reference atlas were also excluded (n=2/26 mice, Extended Data Fig. 8a).

### Linear encoding model

In order to capture the linear contribution of behavioral events to the overall observed neural signal, we fit a linear encoding model for each mouse. The objective is to model the Z-scored photometry signal during the 10 days of defeat with one set of parameters per animal. The feature matrix includes continuous representations of the 4 behaviors of interest quantified by binary random forest classifiers: being attacked, being investigated, fighting back, and fleeing. For the model, the onset and offset of each of the 4 behaviors were considered as 8 behavioral events. To make the contributions of these events temporally flexible, we convolved them with a set of cubic splines^12^. We chose splines spanning from 1 second before to 2 seconds after the behavior to capture both preparatory and reactive neural responses. Then we set the number of splines to 7 based on 3-fold cross-validation accuracy optimized on a linear encoding model. With 7 splines associated with each of the 8 behavior events, we have a total of 56 splines as regressors.

The encoding model takes the form: *y* = β*X* + ε, where *y* is the photometry signal for a given mouse, with dimension *T*, or the number of time points across 10 days. *X* is the design matrix composed of of behavior events convolved with each cubic spline, with dimensions 56x *T*.β is the set of weights for each cubic spline associated with each behavior type, with dimension 1×56. ε is the intercept term.

For every behavior event we calculate the associated kernel as the sum of the 7 splines, each weighted by the learned regression coefficients associated with that behavior event and spline.

### Fast scan cyclic voltammetry

In Extended Data Fig. 5, we recorded fast scan cyclic voltammetry and fiber photometry bulk calcium imaging in the tail of the striatum from a total of 6 sites from 3 male Dat::Cre x Ai148 mice, under a previously published protocol^13^. Mice were injected with urethane (1.5 g/kg dissolved in saline). Once anesthetized, mice were placed within a stereotaxic frame. A stimulating electrode was inserted into the medial forebrain bundle (MFB, AP −2.4 mm, ML +/− 1.1 mm, DV −5 mm from the skull surface at bregma). A carbon-fiber electrode extruding 90-110 μm from a pulled glass capillary was adhered to a 400 μm optical cannula (MFC_400/430-0.48_50mm_MF2.5_FLT from Doric Lenses, Inc.) and inserted into the brain above the tail striatum (AP −1 mm, ML +/− 3.25, DV −2, mm from the brain surface). A silver-chloride-coated reference electrode was inserted nonspecifically in the brain away from the stimulation electrode and recording sites and adhered with a stainless steel skull screw.

To make fiber photometry bulk calcium fluorescence recordings, an optical patch cord was attached to the optic cannula, through which 488 nm excitation laser (400 Hz, 50% duty cycle) delivered excitation, and emission light was received and collected with a photodetector (New Focus, Femtowatt Photoreceiver model 2151). Fluorescence was collected through a usb-powered data acquisition device (USB-201, Measurement Computing) and visualized and stored with TracerDAQ data acquisition software.

The electrode was cycled above the TS by applying a triangular waveform for 10-20 minutes at 60 Hz, followed by 5-10 minutes at 10 Hz. Then the electrode was progressively lowered into the brain while calcium recording took place to identify where the fluorescence levels increased markedly as the electrode entered the striatum. Final recordings were obtained at depths of −2.1 to −2.9 mm below the brain surface.

Upon entering the striatum, voltammetric recordings (LabVIEW software, National Instruments) took place simultaneously with fiber photometry. In each trial, a triangular waveform was applied at 60 Hz to the recording electrode every 100 ms, and after 5s of baseline, the MFB was stimulated with pulses of biphasic stimulation (300 μA, 4ms) after which 25s of recording was collected for a total of 30s per recording. For each recording site, 5 trials for each of 1, 6, 12, 24, and 60 MFB stimulation pulses were delivered. We omit analysis of the 60 pulse trials as the calcium fluorescence signal saturated at 24 pulses.

### Closed-loop, behavior-triggered activation during defeat

To deliver light for optogenetic stimulation during specific behavioral timepoints, we used pre-trained behavior quantification tools described above for inference on video frames streamed in real time.

Images were acquired by a FLIR BlackFlyS camera connected directly to our behavior inference computer (Ubuntu 16.04, equipped with an Nvidia GeForce RTX 2080 Ti graphics card). Using custom code, each video frame was captured by a software trigger (PySpin) and sent as an input to our pre-trained DeepLabCut network for estimating the positions of the interacting mice. The 12 features we defined above were calculated with minor modifications (no smoothing, using adjacent frames for instantaneous speed and velocity features). We trained a separate binary random forest classifier to detect fighting behavior from the unsmoothed features using the same training set as mentioned above for the offline analysis. Upon detection of a fighting back video frame, a serial signal was passed via USB to an arduino, which translated the signal into a TTL for triggering the laser light delivery protocol. The frame capture, behavior inference, and trigger delivery code was run in an open loop and could achieve a speed of ~13 frames per second. A list of timestamps from each frame as well as it’s probability of fight back detection and whether a trigger was delivered were saved for synchronization.

To estimate the latency between behavior detection and light delivery, we measured the differences between the timestamp first real time video frame of a detected fight back bout and the next recorded video frame in which an LED that mirrored laser light was illuminated. This latency estimate may be up to 8 ms (1/120 FPS) slower than the true latency because the LED (and thus laser) may have been illuminated between recorded video camera frames.

Green (532 nm, 0.8 mW for ChRmine animals or 4 mW for NpHR animals) or blue (447 or 460 nm, 7-12 mW for ChR2 animals) lasers were connected to a commutator (Doric Lenses, FRJ_1×2i_FC-2FC_0.22), which led to 200 μM diameter patch cords that were fastened to the mice’s implants via plastic sleeves surrounded by black electric tape. Phasic stimulation was delivered via 3 or 5 pulses of 5 ms in duration.

For fight-back-triggered activation, pulses were continuous so long as fight-back behavior was ongoing to ChRmine animals (compared to being delivered in sets of 5 with a 0.5s pause between sets for ChR2 animals). In all cases, pulses were 5 ms each at 20 Hz. Laser stimulation pulses were recorded for synchronization.

Open-loop activation animals were defeated concurrently with fight-back-triggered animals and received the same light stimulation but tethered to their neighbor’s behavior instead of their own.

Escape-triggered activation animals expressed ChRmine and received a set of 3 green light (0.8 mw) pulses at 20 Hz upon exiting close proximity to the aggressor (10.5 cm) following attack bouts in which fleeing was also detected in at least 50% of attack frames.

Attack-triggered inhibition animals expressed NpHR and received 2s of continuous green light (4 mW) following the onset of an attack.

A separate FLIR BlackFlyS camera was used to record behavior. The output signals from both the real-time inference camera and the recording camera were saved for synchronization.

Across all stimulation paradigms, stimulation was performed across all 10 days of defeat, but not during post-hoc testing.

### Whole brain histology for photometry fiber recovery

Following the completion of our photometry and behavioral studies, animals were deeply anesthetized with an injected ketamine/xylazine cocktail, perfused with 4% PFA dissolved in 1xPBS. Brains were extracted, post-fixed in 4% PFA for 12-24 hours after which they were cleared using the previously described iDISCO+ protocol^64^. Cleared brains were then imaged with light sheet microscopy at 1.1x or 1.3x magnification and those images were registered to a common reference atlas as described previously.^65^ We chose to register to the Paxinos Atlas^66^, in order to access anatomically-defined boundaries between nucleus accumbens core and shell as well as definitions of subregions in the tail of the striatum. Fiber tip locations were manually selected in registered lightsheet images and plotted together in the common Paxinos coordinate frame.

### Histology for optogenetic viral expression and fiber recovery

Following the completion of our optogenetic studies, animals were deeply anesthetized with an injected ketamine/xylazine cocktail, perfused with 4% PFA dissolved in 1x PBS. Brains were extracted, post-fixed in 4% PFA for 12-24 hours after which they were cryoprotected in 30% sucrose and embedded in optimal cutting temperature mounting medium (Fisher Healthcare Tissue-Plus O. C. T. Compound, Fisher Scientific) for freezing over dry ice. Cryosections of the frozen tissue (30 μM slices) were made and stamped directly onto glass microscope slides. Slices were washed with PBS or for immunohistochemistry PBS+0.4% triton (PBST). Then, for immunostaining, blocking buffer (PBST with 2% normal donkey serum and 1% bovine serum albumin) was applied for 30 minutes, followed by incubation by a primary antibody at 4° for 12-24 hours. Following primary incubation, slides were again washed with PBST (3 rounds of 10 minutes each), followed by incubation at room temperature in a secondary antibody for 1 hour, and a final set of washes in PBS (3 rounds of 10 minutes each). Stained or unstained slides were then dried, coverslipped with mounting medium (EMS Immuno Mount DAPI and DABSCO, Electron Microscopy Sciences, Cat # 17989-98, Lot 180418), and sealed with clear nail polish around the slide edges. After at least 12 hours of drying, slides were imaged with a digital robotic slide scanner (NanoZoomer S60, C13210-01, Hamamatsu). Antibodies used: anti-GFP molecular probes G10362, rabbit monoclonal; Alexa fluor 488 Jackson Immuno research 711-545-152.

### Statistical analysis

Sample sizes were not predetermined ahead of time. Experimenters were not blind to animals’ experimental group assignments. The following statistical tests were run in Python version 3.6.12; packages used were scipy version 1.5.2 and statsmodels version 0.12.0.

Tests for linear relationships were Pearson’s correlation (corr, scipy.stats.pearsonr) where repeated measures were not made and general estimated equations (GEE, statsmodels.formula.api.GEE, groups=‘mice’, family=‘Gaussian’, cov_structure=‘Independence) where repeated measures were made. GEEs are extensions to generalized linear models for longitudinal data, where no assumptions are made about the normality of the data and inferences are drawn over the population of mice while also accounting for correlations in measurements within each mouse. One-way ANOVAs (statsmodels.stats.anova.anova_lm), were used when assessing the difference in means across more than two groups, with post-hoc t-Tests (scipy.stats.ttest_ind, equal_var=True, nan_policy=‘propagate’, permutations=None, random_state=None, alternative=‘two-sided’, trim=0) applied where a significant between-group difference was detected. For tests of an interaction between time and more than 2 groups, a 2-way ANOVA (statsmodels.stats.anova.anova_lm, type=2) was used. For tests with more than 2 groups, a mixed linear model (statsmodels.formula.api.mixedlm, fixed effects of endogenous variables of interest, and random effects of intercepts for each mouse) was used. To test equality of variance Levene’s test (scipy.stats.levene, center-median’) was used. Statistical significance in figures and text is set at *p<0.05, **p<0.01, and ***p<0.001.

## Acknowledgements

We thank C. Peña, J. Shaevitz, and the Witten and Falkner laboratories for discussion. We also thank E. Engel for reagents. Funding was from NIH T32MH065214 (L.W.), NSF GRFP DGE-2039656 (L.W.), NIMH DP2MH126375 (A.F.), NIH R01MH126035 (A.F.), and an Alfred P. Sloan Fellowship (A.F.).

## Author contributions

L.W., I.W. and A.F. conceived the project. L.W. and C.C. collected data. L.W. and J.Y. analyzed data. A.F. and I.W. advised on the data analysis. L.W., I.W., and A.F. wrote the paper.

## Code availability

All code will be made available on github (https://github.com/lwillmore/QuantifyingDefeat).

## Data availability

The data that support the findings of this study are available from the corresponding authors upon reasonable request.

