## Supplemental Tables for "Behavioral and dopaminergic signatures of resilience"

**Extended Data Table 1: Statistical significance testing (outside of linear models)**

| FIGURE | Description | Sample size | Statistical test | Statistical values | Test Statistic | *p* Value | Significance |
|---|---|---|---|---|---|---|---|
| Fig. 1n | Cross validated prediction of resilience based on t-SNE behavior | 32 mice | One-sided normal test for proportion > 0.5 | Difference in Proportion | Z=4.529 | p<0.0001 | *** |
| Fig. 2b | TS(DAT::GCaMP) after air puff | 27 mice | Paired t-Test | Difference in Means | t=5.11 | p<0.0001 | *** |
| Fig. 2b | NAc(DAT::GCaMP) after air puff | 27 mice | Paired t-Test | Difference in Means | t=-1.34 | p=0.19 | NS |
| Fig. 2c | TS(DAT::GCaMP) after food approach | 27 mice | Paired t-Test | Difference in Means | t=2.36 | p=0.025 | * |
| Fig. 2c | NAc(DAT::GCaMP) after food approach | 27 mice | Paired t-Test | Difference in Means | t=6.59 | p<0.0001 | *** |
| Fig. 3c | TS(DAT::GCaMP) at social zone entry | 23 mice | Corr | Linear relationship | R=-0.32 | p=0.13 | NS |
| Fig. 3c | TS(DAT::GCaMP) at social zone exit | 23 mice | Corr | Linear relationship | R=0.25 | p=0.23 | NS |
| Fig. 3e | NAc(DAT::GCaMP) at social zone entry | 23 mice | Corr | Linear relationship | R=0.51 | 0.01 | * |
| Fig. 3e | NAc(DAT::GCaMP) at social zone exit | 23 mice | Corr | Linear relationship | R=-0.39 | p=0.062 | NS |
| Fig. 4c | Fight-back onset kernel vs SI time | 19 mice | Corr | Linear relationship | R=0.468 | p=0.043 | * |
| Fig. 4c | Fight-back offset kernel vs SI time | 19 mice | Corr | Linear relationship | R=-0.432 | p=0.065 | NS |
| Fig. 4c | Flee onset kernel vs SI time | 19 mice | Corr | Linear relationship | R=-0.802 | p=0.00004 | *** |
| Fig. 4c | Flee offset kernel vs SI time | 19 mice | Corr | Linear relationship | R=0.58 | p=0.009 | ** |
| Fig. 4c | Being attacked onset kernel vs SI time | 19 mice | Corr | Linear relationship | R=0.51 | p=0.025 | * |
| Fig. 4c | Being attacked offset kernel vs SI time | 19 mice | Corr | Linear relationship | R=-0.603 | p=0.006 | ** |
| Fig. 4c | Being investigated onset kernel vs SI time | 19 mice | Corr | Linear relationship | R=0.28 | p=0.24 | NS |

| | | | | | | | |
|---|---|---|---|---|---|---|---|
| Fig. 4c | Being investigated offset kernel vs SI time | 19 mice | Corr | Linear relationship | R=-0.08 | p=0.74 | NS |
| Fig. 5f | SI time difference across open loop vs no opsin control | 14 mice, 14 mice | t-Test | Difference in mean | t=2.19 | p=0.037 | * |
| Fig. 5g | Sensory period distance difference across open loop vs no opsin control | 14 mice, 14 mice | t-Test | Difference in mean | t=-2.9 | p=0.008 | ** |
| Fig. 5h | Open arm entries difference across open loop vs no opsin control | 14 mice, 14 mice | t-Test | Difference in mean | t=2.56 | p=0.016 | * |
| Fig. 5i | Chamber crossing difference across open loop vs no opsin control | 14 mice, 14 mice | t-Test | Difference in mean | t=2.45 | p=0.02 | * |
| Fig. 5j | Immobility difference across open loop vs no opsin control | 14 mice, 14 mice | t-Test | Difference in mean | t=-1.57 | p=0.12 | NS |
| Fig. 5m | SI time difference across fight-triggered vs no opsin control | 16 mice, 14 mice | t-Test | Difference in mean | t=2.29 | p=0.03 | * |
| Fig. 5n | Sensory period distance difference across fight-triggered vs no opsin control | 16 mice, 14 mice | t-Test | Difference in mean | t=-2.99 | p=0.006 | ** |
| Fig. 5o | Open arm entries difference across fight-triggered vs no opsin control | 16 mice, 14 mice | t-Test | Difference in mean | t=2.87 | p=0.008 | ** |
| Fig. 5p | Chamber crossing difference across fight-triggered vs no opsin control | 16 mice, 14 mice | t-Test | Difference in mean | t=2.64 | p=0.013 | * |
| Fig. 5q | Immobility difference across fight-triggered vs no opsin control | 16 mice, 14 mice | t-Test | Difference in mean | t=-3.05 | p=0.005 | ** |
| Fig. 5t | SI time difference across escape-triggered vs no opsin control | 8 mice, 5 mice | t-Test | Difference in mean | t=0.17 | p=0.87 | NS |
| Fig. 5u | Sensory period distance difference across escape-triggered vs no opsin control | 8 mice, 5 mice | t-Test | Difference in mean | t=-3.94 | p=0.07 | NS |

| | | | | | | | |
|---|---|---|---|---|---|---|---|
| Fig. 5v | Open arm entries difference across escape-triggered vs no opsin control | 8 mice, 5 mice | t-Test | Difference in mean | t=-0.1 | p=0.07 | NS |
| Fig. 5w | Chamber crossing difference across escape-triggered vs no opsin control | 8 mice, 5 mice | t-Test | Difference in mean | t=1.76 | p=0.11 | NS |
| Fig. 5x | Immobility difference across escape-triggered vs no opsin control | 8 mice, 5 mice | t-Test | Difference in mean | t=-0.8 | 0.39 | NS |
| Ext. Fig. 1b | SI ratio difference across stress groups | 54 mice, 3 groups | 1-way ANOVA | Difference in mean | $F_{(51,2)}$=7.015 | p=0.002 | ** |
| Ext. Fig. 1b | SI ratio difference between control and susceptible groups | 22 mice, 13 mice | t-Test | Difference in mean | t=2.793 | p=0.009 | ** |
| Ext. Fig. 1b | SI ratio difference between resilient and susceptible groups | 19 mice, 13 mice | t-Test | Difference in mean | t=3.579 | p=0.001 | ** |
| Ext. Fig. 1b | Object interaction ratio difference across stress groups | 54 mice, 3 groups | 1-way ANOVA | Difference in mean | $F_{(51,2)}$=9.442 | p=3E-4 | *** |
| Ext. Fig. 1b | Object interaction ratio difference between control and susceptible groups | 22 mice, 13 mice | t-Test | Difference in mean | t=-3.185 | p=0.001 | ** |
| Ext. Fig. 1b | Object interaction ratio difference between resilient and susceptible groups | 19 mice, 13 mice | t-Test | Difference in mean | t=-3.185 | p=0.003 | ** |
| Ext. Fig. 1b | SI time difference across stress groups | 54 mice, 3 groups | 1-way ANOVA | Difference in mean | $F_{(51,2)}$=42.9 | p=1E-11 | *** |
| Ext. Fig. 1b | SI time difference between control and susceptible groups | 22 mice, 13 mice | t-Test | Difference in mean | t=8.239 | p=1.6E-9 | *** |
| Ext. Fig. 1b | SI time difference between resilient and susceptible groups | 19 mice, 13 mice | t-Test | Difference in mean | t=8.547 | p=1.55E-9 | *** |
| Ext. Fig. 1b | Object interaction time difference across stress groups | 54 mice, 3 groups | 1-way ANOVA | Difference in mean | $F_{(51,2)}$=27.26 | p=9E-9 | *** |

| | | | | | | | |
|---|---|---|---|---|---|---|---|
| Ext. Fig. 1b | Object interaction time difference between control and susceptible groups | 22 mice, 13 mice | t-Test | Difference in mean | t=-6.08 | p=7.62E-7 | *** |
| Ext. Fig. 1b | Object interaction time difference between resilient and susceptible groups | 19 mice, 13 mice | t-Test | Difference in mean | t=-5.82 | p=2.3E-8 | *** |
| Ext. Fig. 1c | Sucrose preference difference between stress groups | 54 mice, 3 groups | 1-way ANOVA | Difference in mean | $F_{(51,2)}$=3.21 | p=0.048 | * |
| Ext. Fig. 1c | Sucrose preference difference between control and susceptible groups | 22 mice, 13 mice | t-Test | Difference in mean | t=2.15 | p=0.039 | * |
| Ext. Fig.1c | Sucrose preference difference between resilient and susceptible groups | 19 mice, 13 mice | t-Test | Difference in mean | t=2.37 | p=0.024 | * |
| Ext. Fig. 1d | SI time vs sucrose preference | 54 mice | Pearson's correlation | Linear relationship | R=0.294 | p=0.03 | * |
| Ext. Fig. 1e | Difference in weight change on every CSDS day between resilient and susceptible mice and between control and susceptible mice | 54 mice, 3 groups, 10 days | t-Tests, 20 comparisons for FDR correction | Difference in mean | - | p<0.05 shown | * |
| Ext. Fig. 1e | Absolute body weight on every CSDS day between resilient and susceptible mice and between control and susceptible mice | 54 mice, 3 groups, 10 days | t-Tests, 20 comparisons for FDR correction | Difference in mean | - | p<0.05 shown | * |
| Ext. Fig. 1g | Crossings difference between stress groups | 54 mice, 3 groups | 1-way ANOVA | Difference in mean | $F_{(51,2)}$=5.77 | p=0.006 | ** |
| Ext. Fig. 1g | Crossings difference between control and susceptible groups | 22 mice, 13 mice | t-Test | Difference in mean | t=2.69 | p=0.01 | * |
| Ext. Fig. 1g | Crossings difference between resilient and susceptible groups | 19 mice, 13 mice | t-Test | Difference in mean | t=2.78 | p=0.008 | ** |

| | | | | | | | |
|---|---|---|---|---|---|---|---|
| Ext. Fig. 1h | Immobility difference between stress groups | 54 mice, 3 groups | 1-way ANOVA | Difference in mean | $F_{(51,2)}$=3.16 | p=0.05 | * |
| Ext. Fig. 1h | Immobility difference between control and susceptible groups | 22 mice, 13 mice | t-Test | Difference in mean | t=-2.24 | p=0.03 | * |
| Ext. Fig. 1h | Immobility difference between resilient and susceptible groups | 19 mice, 13 mice | t-Test | Difference in mean | t=-1.86 | p=0.07 | NS |
| Ext. Fig. 1j | SI time vs dominance | 54 mice | Pearson's correlation | Linear relationship | R=-0.038 | p=0.78 | NS |
| Ext. Fig. 1k | Dominance difference between stress groups | 54 mice, 3 groups | 1-way ANOVA | Difference in mean | $F_{(51,2)}$=0.34 | p=0.71 | NS |
| Ext. Fig. 1l | Time spent interacting with freely moving male target difference between stress groups | 35 mice, 3 groups, 2 times | 2-way ANOVA | Interaction of factors | $F_{(64,2)}$=0.31 | p=0.72 | NS |
| Ext. Fig. 1m | Time spent interacting with freely moving female target difference between stress groups | 35 mice, 3 groups, 2 times | 2-way ANOVA | Interaction of factors | $F_{(64,2)}$=0.189 | p=0.828 | NS |
| Ext. Fig 3b | Average t-SNE behavior cluster occupancy difference between resilient and susceptible groups | 32 mice | t-Tests | Difference in mean | - | p<0.05 shown | * |
| Ext. Fig 3b | Average t-SNE behavior cluster occupancy difference in variance between resilient and susceptible groups | 32 mice | Levene's tests | Difference in variance | - | p<0.05 shown | * |
| Ext. Fig. 4f | Fighting while attacked difference between males and females | 32 males, 8 females | t-Tests | Difference in mean | t=-7.45 | p<0.001 | *** |
| Ext. Fig. 7b | TS(DAT::GCaMP) at social zone entry | 23 mice | Corr | Linear relationship | R=-0.32 | p=0.13 | NS |
| Ext. Fig. 7b | TS(DAT::GCaMP) at social zone exit | 23 mice | Corr | Linear relationship | R=0.25 | p=0.23 | NS |
| Ext. Fig. 7d | TS(DAT::GCaMP) at object zone entry | 23 mice | Pearson's correlation | Linear relationship | R=0.06 | p=0.8 | NS |
| Ext. Fig. 7d | TS(DAT::GCaMP) at object zone exit | 23 mice | Pearson's correlation | Linear relationship | R=0.34 | p=0.11 | NS |

| | | | | | | | |
|---|---|---|---|---|---|---|---|
| Ext. Fig. 7f | NAc(DAT::GCaMP) at object zone entry | 23 mice | Pearson's correlation | Linear relationship | R=0.15 | p=0.47 | NS |
| Ext. Fig. 7f | NAc(DAT::GCaMP) at object zone exit | 23 mice | Pearson's correlation | Linear relationship | R=0.15 | p=0.47 | NS |
| Ext. Fig. 8a | Kernel weight at behavior vs % behavior expression | 19 mice | Pearson's correlation | Linear relationship | - | p<0.05 shown | * |
| Ext. Fig. 8d | Kernel weights at different fiber locations | 19 mice | t-Tests | Difference in mean | - | p<0.05 shown | * |
| Ext. Fig. 8e | Kernel weight from TS encoding model difference between resilient and susceptible mice | 19 mice | t-Tests | Difference in mean | - | p>0.05 | NS |
| Ext. Fig. 9g | Fighting while attacked early vs late in open loop | 14 mice | Paired t-Test | Difference in mean | t=0.38 | p=0.71 | NS |
| Ext. Fig. 9g | Fighting while attacked early vs late in fight-triggered | 16 mice | Paired t-Test | Difference in mean | t=2.77 | p=0.04 | * |
| Ext. Fig. 9g | Fighting while attacked early vs late in fight-triggered control | 14 mice | Paired t-Test | Difference in mean | t=-0.41 | p=0.71 | NS |
| Ext. Fig. 9g | Fighting while attacked early vs late in escape-triggered | 8 mice | Paired t-Test | Difference in mean | t=-2.33 | p=0.1 | NS |
| Ext. Fig. 9g | Fighting while attacked early vs late in escape-triggered control | 5 mice | Paired t-Test | Difference in mean | t=-1.14 | p=0.34 | NS |
| Ext. Fig. 9h | Fleeing while attacked early vs late in open loop | 14 mice | Paired t-Test | Difference in mean | t=1.12 | p=0.42 | NS |
| Ext. Fig. 9h | Fleeing while attacked early vs late in fight-triggered | 16 mice | Paired t-Test | Difference in mean | t=-0.29 | p=0.78 | NS |
| Ext. Fig. 9h | Fleeing while attacked early vs late in fight-triggered control | 14 mice | Paired t-Test | Difference in mean | t=-1.49 | p=0.42 | NS |

| | | | | | | | |
|---|---|---|---|---|---|---|---|
| Ext. Fig. 9h | Fleeing while attacked early vs late in escape-triggered | 8 mice | Paired t-Test | Difference in mean | t=6.42 | p<0.001 | *** |
| Ext. Fig. 9h | Fleeing while attacked early vs late in escape-triggered control | 5 mice | Paired t-Test | Difference in mean | t=0.68 | p=0.54 | NS |
| Ext. Fig. 9o | Time being attacked inhibition vs control | 12 mice, 10 mice | t-Test | Difference in mean | t=-3.03 | p=0.007 | ** |
| Ext. Fig. 9p | Time fleeing inhibition vs control | 12 mice, 10 mice | t-Test | Difference in mean | t=-3.92 | p<0.001 | *** |
| Ext. Fig. 9q | SI time difference across attack-triggered inhibition vs no opsin control | 12 mice, 10 mice | t-Test | Difference in mean | t=0.04 | p=0.96 | NS |
| Ext. Fig. 9r | Sensory period distance difference across attack-triggered inhibition vs no opsin control | 12 mice, 10 mice | Levene Test | Difference in variance | W=4.92 | p=0.04 | * |
| Ext. Fig. 9s | Open arm entries difference across attack-triggered inhibition vs no opsin control | 12 mice, 10 mice | t-Test | Difference in mean | t=-0.64 | p=0.53 | NS |
| Ext. Fig. 9t | Chamber crossing difference across attack-triggered inhibition vs no opsin control | 12 mice, 10 mice | t-Test | Difference in mean | t=-0.024 | p=0.98 | NS |
| Ext. Fig. 9u | Immobility difference attack-triggered inhibition vs no opsin control | 12 mice, 10 mice | t-Test | Difference in mean | t=-0.47 | p=0.64 | NS |

| *Model: SI time = weight change + intercept, grouped by mouse* | **$\beta$±standard error** | **z-stat** | **p-value** | **95% CI [lower, upper]** | |
|---|---|---|---|---|---|
| **Intercept** | 39.426±3.085 | 12.779 | <0.001 | 33.379 | 45.474 |
| **SI time** | 0.740±0.369 | 2.007 | 0.045 | 0.017 | 1.462 |

**Extended Data Table 2:** GEE regression (Extended Data Fig. 1e): *SI time = weight change from day 1 + intercept, grouped by mouse*. Number of mice (groups) = 32, minimum samples per group 8, maximum samples per group 9, dependence structure = independence, family = Gaussian.

| *Model: % time being attacked = SI time + intercept, grouped by mouse* | **β±standard error** | **z-stat** | **p-value** | **95% CI [lower, upper]** | |
|---|---|---|---|---|---|
| **Intercept** | 12.086±0.986 | 12.258 | <0.001 | 10.153 | 14.018 |
| **SI time** | -0.003±0.026 | -0.114 | 0.909 | -0.053 | 0.047 |

**Extended Data Table 3:** GEE regression (Fig. 1i): *% time being attacked = (SI time + intercept, grouped by mouse*. Number of mice (groups) = 32, minimum samples per group 9, maximum samples per group 10, dependence structure = independence, family = Gaussian.

| *Model: % time being investigated = SI time + intercept, grouped by mouse* | **β±standard error** | **z-stat** | **p-value** | **95% CI [lower, upper]** | |
|---|---|---|---|---|---|
| **Intercept** | 11.055±1.383 | 7.992 | <0.001 | 8.344 | 13.766 |
| **SI time** | -0.067±0.033 | -2.052 | 0.040 | -0.132 | -0.003 |

**Extended Data Table 4:** GEE regression (Fig. 1i): *% time being investigated = (SI time + intercept, grouped by mouse*. Number of mice (groups) = 32, minimum samples per group 9, maximum samples per group 10, dependence structure = independence, family = Gaussian.

| *Model: % fighting back while attacked = SI time + intercept, grouped by mouse* | **β±standard error** | **z-stat** | **p-value** | **95% CI [lower, upper]** | |
|---|---|---|---|---|---|
| **Intercept** | 21.116±2.361 | 8.943 | <0.001 | 16.488 | 25.743 |
| **SI time** | 0.1734±0.072 | 2.419 | 0.016 | 0.033 | 0.314 |

**Extended Data Table 5:** GEE regression (Fig. 1i): *% time fighting back while attacked = (SI time + intercept, grouped by mouse*. Number of mice (groups) = 32, minimum samples per group 9, maximum samples per group 10, dependence structure = independence, family = Gaussian.

| *Model: % fleeing back while attacked = SI time + intercept, grouped by mouse* | **β±standard error** | **z-stat** | **p-value** | **95% CI [lower, upper]** | |
|---|---|---|---|---|---|
| **Intercept** | 69.747±4.069 | 17.140 | <0.001 | 61.771 | 77.723 |
| **SI time** | -0.1851±0.114 | -1.626 | 0.104 | -0.408 | 0.038 |

**Extended Data Table 6:** GEE regression (Fig. 1i): *% time fleeing while attacked = (SI time + intercept, grouped by mouse*. Number of mice (groups) = 32, minimum samples per group 9, maximum samples per group 10, dependence structure = independence, family = Gaussian.

| *Model: % time males are attacked = defeat_day + resilience_category + defeat_day*resilience_category + intercept* | **β±standard error** | **z-stat** | **p-value** | **95% CI [lower, upper]** | |
|---|---|---|---|---|---|
| **Intercept** | 12.7654±1.596 | 7.998 | <0.001 | 9.637 | 15.893 |
| **defeat day** | -0.170±0.277 | -0.613 | 0.54 | -0.712 | 0.373 |

| | | | | | |
|---|---|---|---|---|---|
| **resilience category** | 0.505±1.879 | 0.269 | 0.788 | -3.178 | 4.188 |
| **defeat day * resilience category** | -0.0026±0.013 | -0.194 | 0.846 | -0.029 | 0.024 |

**Extended Data Table 7:** GEE regression of behavior (Extended Data Fig. 2): *% time males are attacked = defeat_day + resilience_category + defeat_day*resilience_category + intercept, grouped by mouse*. Number of mice (groups) = 32, minimum samples per group 8, maximum samples per group 9, dependence structure = independence, family = Gaussian.

| *Model: % time males are investigated = defeat_day + resilience_category + defeat_day*resilience_category + intercept* | **β±standard error** | **z-stat** | **p-value** | **95% CI [lower, upper]** | |
|---|---|---|---|---|---|
| **Intercept** | 5.421±0.920 | 5.895 | <0.001 | 3.619 | 7.223 |
| **defeat day** | 0.581±0.145 | 4.005 | <0.001 | 0.296 | 0.865 |
| **resilience category** | -0.039±1.25 | -0.031 | 0.975 | -2.489 | 2.411 |
| **defeat day * resilience category** | -0.0913±0.236 | -0.387 | 0.699 | -0.554 | 0.371 |

**Extended Data Table 8:** GEE regression of behavior (Extended Data Fig. 2): *% time males are investigated = defeat_day + resilience_category + defeat_day*resilience_category + intercept, grouped by mouse*. Number of mice (groups) = 32, minimum samples per group 8, maximum samples per group 9, dependence structure = independence, family = Gaussian.

| *Model: % males fight back while attacked = defeat_day + resilience_category + defeat_day*resilience_category + intercept* | **β±standard error** | **z-stat** | **p-value** | **95% CI [lower, upper]** | |
|---|---|---|---|---|---|
| **Intercept** | 25.368±2.692 | 9.423 | <0.001 | 20.092 | 30.644 |
| **defeat day** | 0.001±0.498 | 0.020 | 0.984 | -0.966 | 0.986 |
| **resilience category** | 3.006±3.335 | 0.901 | 0.367 | -3.531 | 9.543 |
| **defeat day * resilience category** | 0.308±0.551 | 0.559 | 0.576 | -0.772 | 1.388 |

**Extended Data Table 9:** GEE regression of behavior (Extended Data Fig. 2): *% males fight back while attacked = defeat_day + resilience_category + defeat_day*resilience_category + intercept, grouped by mouse*. Number of mice (groups) = 32, minimum samples per group 8, maximum samples per group 9, dependence structure = independence, family = Gaussian.

| *Model: % males flee while attacked = defeat_day + resilience_category + defeat_day*resilience_category + intercept* | **β±standard error** | **z-stat** | **p-value** | **95% CI [lower, upper]** | |
|---|---|---|---|---|---|
| **Intercept** | 60.715±3.468 | 17.507 | <0.001 | 53.918 | 67.513 |
| **defeat day** | 0.782±0.444 | 1.762 | 0.078 | -0.088 | 1.652 |
| **resilience category** | 0.898±4.305 | 0.209 | 0.835 | -7.540 | 9.336 |
| **defeat day * resilience category** | -1.0337±0.558 | -1.854 | 0.064 | -2.127 | 0.059 |

**Extended Data Table 10:** GEE regression of behavior (Extended Data Fig. 2): *% males flee while attacked = defeat_day + resilience_category + defeat_day*resilience_category + intercept, grouped by mouse*. Number of mice (groups) = 32, minimum samples per group 8, maximum samples per group 9, dependence structure = independence, family = Gaussian.

| *Model: % time females are attacked = SI time + intercept, grouped by mouse* | **β±standard error** | **z-stat** | **p-value** | **95% CI [lower, upper]** | |
|---|---|---|---|---|---|
| **Intercept** | 13.6884±1.536 | 8.912 | <0.001 | 10.678 | 16.699 |
| **SI time** | -0.0122±0.04 | -0.308 | 0.758 | -0.09 | 0.066 |

**Extended Data Table 3:** GEE regression (Extended Data Fig. 4f): *% time females are attacked = SI time + intercept, grouped by mouse*. Number of mice (groups) = 8, minimum samples per group 10, maximum samples per group 10, dependence structure = independence, family = Gaussian.

| *Model: % time females are investigated = SI time + intercept, grouped by mouse* | **β±standard error** | **z-stat** | **p-value** | **95% CI [lower, upper]** | |
|---|---|---|---|---|---|
| **Intercept** | 12.0307±2.698 | 4.459 | <0.001 | 6.743 | 17.319 |
| **SI time** | 0.052±0.046 | 1.121 | 0.262 | -0.039 | 0.143 |

**Extended Data Table 4:** GEE regression (Extended Data Fig. 4f): *% time females are investigated = SI time + intercept, grouped by mouse*. Number of mice (groups) = 8, minimum samples per group 10, maximum samples per group 10, dependence structure = independence, family = Gaussian.

| *Model: % time females fight back while attacked = SI time + intercept, grouped by mouse* | **β±standard error** | **z-stat** | **p-value** | **95% CI [lower, upper]** | |
|---|---|---|---|---|---|
| **Intercept** | 12.399±1.71 | 7.2493 | <0.001 | 9.047 | 15.751 |
| **SI time** | -0.0358±0.025 | -1.42 | 0.155 | -0.085 | 0.014 |

**Extended Data Table 5:** GEE regression (Extended Data Fig. 4f): *% females fight back while attacked = SI time + intercept, grouped by mouse*. Number of mice (groups) = 8, minimum samples per group 10, maximum samples per group 10, dependence structure = independence, family = Gaussian.

| *Model: % time females flee while attacked = SI time + intercept, grouped by mouse* | **β±standard error** | **z-stat** | **p-value** | **95% CI [lower, upper]** | |
|---|---|---|---|---|---|
| **Intercept** | 62.3257±2.503 | 24.903 | <0.001 | 57.420 | 67.231 |
| **SI time** | 0.0420±0.033 | 1.279 | 0.201 | -0.022 | 0.106 |

**Extended Data Table 6:** GEE regression (Extended Data Fig. 4f): *% females flee while attacked = SI time + intercept, grouped by mouse*. Number of mice (groups) = 8, minimum samples per group 10, maximum samples per group 10, dependence structure = independence, family = Gaussian.

| *Model: % time females spent in cluster 5 = SI time + intercept, grouped by mouse* | **β±standard error** | **z-stat** | **p-value** | **95% CI [lower, upper]** | |
|---|---|---|---|---|---|
| **Intercept** | 6.71±1.02 | 6.541 | <0.001 | 4.7 | 8.721 |
| **SI time** | 0.086±0.019 | 4.575 | <0.001 | 0.049 | 0.123 |

**Extended Data Table 15:** GEE regression (Fig. 1u): *% time females spent in cluster 5 = (SI time + intercept, grouped by mouse*. Number of mice (groups) = 8, minimum samples per group 10, maximum samples per group 10, dependence structure = independence, family = Gaussian.

| *Model:* ***onset*** *Z-score TS(Dat::GCaMP) ΔF/F=defeat_day + SI_time + defeat_day*SI_time + intercept* | **β±standard error** | **z-stat** | **p-value** | **95% CI [upper, lower]** | |
|---|---|---|---|---|---|
| **Intercept** | 0.347±0.099 | 3.491 | <0.001 | 0.152 | 0.542 |
| **defeat day** | 0.0253±0.01 | 2.584 | 0.01 | 0.006 | 0.044 |
| **SI time** | -0.0009±0.002 | -0.473 | 0.636 | -0.005 | 0.003 |
| **defeat day * SI time** | 0.00001±0 | 0.345 | 0.73 | 0 | 0.001 |

**Extended Data Table 16:** GEE regression of TS(Dat::GCaMP) (Fig. 3b,c): *Z-score TS(Dat::GCaMP) ΔF/F at proximity onset=defeat_day + SI_time + defeat_day*SI_time + intercept, grouped by mouse*. Number of mice (groups) = 27, minimum samples per group 8, maximum samples per group 10, dependence structure = independence, family = Gaussian.

| *Model:* ***offset*** *Z-score TS(Dat::GCaMP) ΔF/F=defeat_day + SI_time + defeat_day*SI_time + intercept* | **β±standard error** | **z-stat** | **p-value** | **95% CI [lower, upper]** | |
|---|---|---|---|---|---|
| **Intercept** | -0.2992±0.053 | -5.665 | <0.001 | -0.403 | -0.196 |
| **defeat day** | -0.0057±0.01 | -0.576 | 0.565 | -0.025 | 0.014 |
| **SI time** | -0.00003±0.001 | -0.032 | 0.975 | -0.002 | 0.002 |
| **defeat day * SI time** | 0.0001±0 | 0.555 | 0.579 | 0 | 0.001 |

**Extended Data Table 17:** GEE regression of TS(Dat::GCaMP) (Fig. 3b,c): *Z-score TS(Dat::GCaMP) ΔF/F at proximity offset=defeat_day + SI_time + defeat_day*SI_time + intercept, grouped by mouse*. Number of mice (groups) = 27, minimum samples per group 8, maximum samples per group 10, dependence structure = independence, family = Gaussian.

| *Model: in males,* ***onset*** *Z-score TS(Dat::GCaMP) ΔF/F=defeat_day + resilience_category + defeat_day*resilience_category + intercept* | **β±standard error** | **z-stat** | **p-value** | **95% CI [upper, lower]** | |
|---|---|---|---|---|---|
| **Intercept** | 0.400±0.092 | 4.363 | <0.001 | 0.220 | 0.579 |
| **defeat day** | 0.033±0.006 | 5.710 | <0.001 | 0.022 | 0.045 |
| **resilience category** | -0.131±0.136 | -0.960 | 0.337 | -0.397 | 0.136 |
| **defeat day * resilience category** | 0.008±0.013 | 0.602 | 0.547 | -0.018 | 0.033 |

**Extended Data Table 18:** GEE regression of TS(Dat::GCaMP) (Extended Data Fig. 6b): *Z-score TS(Dat::GCaMP) ΔF/F at proximity onset=defeat_day + resilience_category + defeat_day*resilience_category + intercept, grouped by mouse*. Number of mice (groups) = 19, minimum samples per group 8, maximum samples per group 10, dependence structure = independence, family = Gaussian.

| *Model: in males,* ***offset*** *Z-score TS(Dat::GCaMP) ΔF/F=defeat_day + resilience_category + defeat_day*resilience_category + intercept* | **β±standard error** | **z-stat** | **p-value** | **95% CI [upper, lower]** | |
|---|---|---|---|---|---|
| **Intercept** | -0.351±0.066 | -5.329 | <0.001 | -0.480 | -0.222 |
| **defeat day** | 0.003±0.007 | 0.450 | 0.653 | -0.01 | 0.017 |
| **resilience category** | 0.076±0.082 | 0.924 | 0.355 | -0.085 | 0.237 |
| **defeat day * resilience category** | -0.012±0.009 | -1.365 | 0.172 | -0.03 | 0.005 |

**Extended Data Table 19:** GEE regression of TS(Dat::GCaMP) (Extended Data Fig. 6b): *Z-score NAc(Dat::GCaMP) ΔF/F at proximity offset=defeat_day + resilience_category + defeat_day*resilience_category + intercept, grouped by mouse*. Number of mice (groups) = 19, minimum samples per group 8, maximum samples per group 10, dependence structure = independence, family = Gaussian.

| *Model: in females,* ***onset*** *Z-score TS(Dat::GCaMP) ΔF/F=defeat_day + resilience_category + defeat_day*resilience_category + intercept* | **β±standard error** | **z-stat** | **p-value** | **95% CI [upper, lower]** | |
|---|---|---|---|---|---|
| **Intercept** | 0.4203±0.107 | 3.919 | <0.001 | 0.210 | 0.631 |
| **defeat day** | 0.0083±0.013 | 0.673 | 0.501 | -0.017 | 0.034 |
| **resilience category** | -0.1062±0.111 | -0.955 | 0.340 | -0.324 | 0.112 |
| **defeat day * resilience category** | -0.0029±0.016 | -0.180 | 0.857 | -0.034 | 0.029 |

**Extended Data Table 20:** GEE regression of TS(Dat::GCaMP) (Extended Data Fig. 6b): *Z-score TS(Dat::GCaMP) ΔF/F at proximity onset=defeat_day + resilience_category + defeat_day*resilience_category + intercept, grouped by mouse*. Number of mice (groups) = 8, minimum samples per group 8, maximum samples per group 10, dependence structure = independence, family = Gaussian.

| *Model: in females,* ***offset*** *Z-score TS(Dat::GCaMP) ΔF/F=defeat_day + resilience_category + defeat_day*resilience_category + intercept* | **β±standard error** | **z-stat** | **p-value** | **95% CI [upper, lower]** | |
|---|---|---|---|---|---|
| **Intercept** | -0.2612±0.057 | -4.585 | <0.001 | -0.373 | -0.150 |
| **defeat day** | -0.0088±0.008 | -1.081 | 0.280 | -0.025 | 0.007 |
| **resilience category** | 0.0369±0.058 | 0.639 | 0.523 | -0.076 | 0.150 |
| **defeat day * resilience category** | 0.0238±0.010 | 2.326 | 0.02 | 0.004 | 0.0044 |

**Extended Data Table 21:** GEE regression of TS(Dat::GCaMP) (Extended Data Fig. 6b): *Z-score NAc(Dat::GCaMP) ΔF/F at proximity offset=defeat_day + resilience_category + defeat_day*resilience_category + intercept, grouped by mouse*. Number of mice (groups) = 8, minimum samples per group 8, maximum samples per group 10, dependence structure = independence, family = Gaussian.

| *Model: **onset** Z-score NAc(Dat::GCaMP) ΔF/F=defeat_day + SI_time + defeat_day*SI_time + intercept* | **β±standard error** | **z-stat** | **p-value** | **95% CI [lower, upper]** | |
|---|---|---|---|---|---|
| **Intercept** | -0.3493±0.073 | -4.788 | <0.001 | -0.492 | -0.206 |
| **defeat day** | -0.0012±0.002 | -0.107 | 0.915 | -0.024 | 0.021 |
| **SI time** | 0.0038±0.011 | 1.982 | 0.047 | 0 | 0.008 |
| **defeat day * SI time** | -0.0001±0 | -0.29 | 0.772 | -0.001 | 0.001 |

**Extended Data Table 22:** GEE regression of NAc(Dat::GCaMP) (Fig. 3e,f): *Z-score NAc(Dat::GCaMP) ΔF/F at proximity onset=defeat_day + SI_time + defeat_day*SI_time + intercept, grouped by mouse*. Number of mice (groups) = 27, minimum samples per group 8, maximum samples per group 10, dependence structure = independence, family = Gaussian.

| *Model: **offset** Z-score NAc(Dat::GCaMP) ΔF/F=defeat_day + SI_time + defeat_day*SI_time + intercept* | **β±standard error** | **z-stat** | **p-value** | **95% CI [lower, upper]** | |
|---|---|---|---|---|---|
| **Intercept** | 0.292±0.092 | 3.163 | 0.002 | 0.111 | 0.472 |
| **defeat day** | 0.0047±0.015 | 0.316 | 0.752 | -0.025 | 0.034 |
| **SI time** | -0.004±0.002 | -2.249 | 0.025 | -0.008 | -0.001 |
| **defeat day * SI time** | 0.0004±0 | 1.199 | 0.231 | 0 | 0.001 |

**Extended Data Table 23:** GEE regression of NAc(Dat::GCaMP) (Fig. 3e,f): *Z-score NAc(Dat::GCaMP) ΔF/F at proximity offset=defeat_day + SI_time + defeat_day*SI_time + intercept, grouped by mouse*. Number of mice (groups) = 27, minimum samples per group 8, maximum samples per group 10, dependence structure = independence, family = Gaussian.

| *Model: in males, **onset** Z-score NAc(Dat::GCaMP) ΔF/F=defeat_day + resilience_category + defeat_day*resilience_category + intercept* | **β±standard error** | **z-stat** | **p-value** | **95% CI [upper, lower]** | |
|---|---|---|---|---|---|
| **Intercept** | -0.2553±0.037 | -6.944 | <0.001 | -0.327 | -0.183 |
| **defeat day** | -0.0077±0.006 | -1.328 | 0.184 | -0.019 | 0.004 |
| **resilience category** | 0.1761±0.072 | 2.446 | 0.014 | 0.035 | 0.317 |
| **defeat day * resilience category** | -0.0026±0.013 | -0.194 | 0.846 | -0.029 | 0.024 |

**Extended Data Table 24:** GEE regression of NAc(Dat::GCaMP) (Extended Data Fig. 6d): *Z-score NAc(Dat::GCaMP) ΔF/F at proximity onset=defeat_day + resilience_category + defeat_day*resilience_category + intercept, grouped by mouse*. Number of mice (groups) = 19, minimum samples per group 8, maximum samples per group 10, dependence structure = independence, family = Gaussian.

| *Model: in males, **offset** Z-score NAc(Dat::GCaMP) ΔF/F=defeat_day +* | **β±standard error** | **z-stat** | **p-value** | **95% CI [upper, lower]** |
|---|---|---|---|---|

| resilience_category + defeat_day*resilience_category + intercept | | | | | |
|---|---|---|---|---|---|
| **Intercept** | 0.1407±0.053 | 2.642 | 0.008 | 0.036 | 0.245 |
| **defeat day** | 0.0250±0.005 | 4.966 | <0.001 | 0.015 | 0.035 |
| **resilience category** | -0.1037±0.077 | -1.340 | 0.180 | -0.255 | 0.048 |
| **defeat day * resilience category** | 0.0023±0.009 | 0.262 | 0.793 | -0.015 | 0.020 |

**Extended Data Table 25:** GEE regression of NAc(Dat::GCaMP) (Extended Data Fig. 6d): *Z-score NAc(Dat::GCaMP) ΔF/F at proximity offset=defeat_day + resilience_category + defeat_day*resilience_category + intercept, grouped by mouse*. Number of mice (groups) = 19, minimum samples per group 8, maximum samples per group 10, dependence structure = independence, family = Gaussian.

| *Model: in females,* ***onset*** *Z-score NAc(Dat::GCaMP) ΔF/F=defeat_day + resilience_category + defeat_day*resilience_category + intercept* | **β±standard error** | **z-stat** | **p-value** | **95% CI [upper, lower]** | |
|---|---|---|---|---|---|
| **Intercept** | -0.3367±0.062 | -5.423 | <0.001 | -0.458 | -0.215 |
| **defeat day** | 0.0114±0.008 | 1.369 | 0.171 | -0.005 | 0.028 |
| **resilience category** | 0.1285±0.081 | 1.578 | 0.114 | -0.031 | 0.3288 |
| **defeat day * resilience category** | -0.0136±0.016 | -0.844 | 0.399 | -0.045 | 0.018 |

**Extended Data Table 26:** GEE regression of NAc(Dat::GCaMP) (Extended Data Fig. 6d): *Z-score NAc(Dat::GCaMP) ΔF/F at proximity onset=defeat_day + resilience_category + defeat_day*resilience_category + intercept, grouped by mouse*. Number of mice (groups) = 8, minimum samples per group 8, maximum samples per group 10, dependence structure = independence, family = Gaussian.

| *Model: in females,* ***offset*** *Z-score NAc(Dat::GCaMP) ΔF/F=defeat_day + resilience_category + defeat_day*resilience_category + intercept* | **β±standard error** | **z-stat** | **p-value** | **95% CI [upper, lower]** | |
|---|---|---|---|---|---|
| **Intercept** | 0.2691±0.098 | 2.752 | 0.006 | 0.077 | 0.461 |
| **defeat day** | 0.0008±0.021 | 0.041 | 0.968 | -0.036 | 0.064 |
| **resilience category** | -0.1956±0.121 | -1.621 | 0.105 | -0.432 | 0.041 |
| **defeat day * resilience category** | 0.0138±0.026 | 0.541 | 0.588 | -0.036 | 0.064 |

**Extended Data Table 27:** GEE regression of NAc(Dat::GCaMP) (Extended Data Fig. 6d): *Z-score NAc(Dat::GCaMP) ΔF/F at proximity offset=defeat_day + resilience_category + defeat_day*resilience_category + intercept, grouped by mouse*. Number of mice (groups) = 8, minimum samples per group 8, maximum samples per group 10, dependence structure = independence, family = Gaussian.

| *Model: ΔAOC NAc =defeat_day + resilience_category + defeat_day*resilience_category + intercept* | **β±standard error** | **z-stat** | **p-value** | **95% CI [upper, lower]** |
|---|---|---|---|---|

| | | | | | |
|---|---|---|---|---|---|
| **Intercept** | -6.976±3.889 | -1.794 | 0.073 | -14.599 | 0.647 |
| **defeat day** | 0.711±0.462 | 1.539 | 0.124 | -0.194 | 1.616 |
| **resilience category** | 2.583±4.89 | 0.528 | 0.597 | -7.002 | 12.168 |
| **defeat day * resilience category** | -1.175±0.577 | -2.037 | 0.042 | -2.305 | -0.044 |

**Extended Data Table 28:** Mixed linear regression (Extended Data Fig. 10c): *ΔAOC NAc = defeat_day + resilience_category + defeat_day*resilience_category + intercept, grouped by mouse*. Number of mice (groups) = 19, minimum samples per group 7, maximum samples per group 10, dependence structure = independence, family = Gaussian.

| *Model: ΔAOC TS =defeat_day + resilience_category + defeat_day*resilience_category + intercept* | **β±standard error** | **z-stat** | **p-value** | **95% CI [upper, lower]** | |
|---|---|---|---|---|---|
| **Intercept** | -10.131±9.589 | -1.057 | 0.291 | -28.925 | 8.662 |
| **defeat day** | -1.058±0.927 | -1.141 | 0.254 | -2.875 | 0.76 |
| **resilience category** | -10.358±12.08 | -0.857 | 0.391 | -34.03 | 13.319 |
| **defeat day * resilience category** | 1.215±0.927 | 1.044 | 0.296 | -1.066 | 3.495 |

**Extended Data Table 29:** Mixed linear regression (Extended Data Fig. 10d): *ΔAOC TS = defeat_day + resilience_category + defeat_day*resilience_category + intercept, grouped by mouse*. Number of mice (groups) = 19, minimum samples per group 7, maximum samples per group 10, dependence structure = independence, family = Gaussian.

**Extended Data Table 30: Fiber placement aligned to Paxinos Atlas reference coordinates**

| mouse | SI time (%) | NAc region | NAc AP (cm) | NAc ML (cm) | NAc DV (cm) | TS region (cm) | TS AP (cm) | TS ML (cm) | TS DV (cm) |
|---|---|---|---|---|---|---|---|---|---|
| 355 | 55.211 | core | 1 | 1.06 | -5.81 | intermediate/dorsal | -0.8 | -2.98 | -3.55 |
| 353 | 36.548 | core | 1.3 | -1.05 | -5.18 | intermediate/dorsal | -0.8 | 3.06 | -3.51 |
| 337 | 29.2963 | core | 1.1 | 1.31 | -5.86 | intermediate/dorsal | -1 | -3.07 | -3.42 |
| 332 | 55.1 | core | 0.9 | 0.97 | -5.44 | intermediate/dorsal | -1.3 | -3.46 | -3.46 |
| 335 | 27.2609333 | shell | 0.7 | 0.78 | -5.89 | intermediate/dorsal | -1.1 | -2.96 | -3.77 |
| 346 | 32.0212333 | core | 0.4 | -1.08 | -5.97 | intermediate | -1.5 | 3.23 | -3.69 |
| 37 | 54.555 | shell | 1 | 0.79 | -6.07 | intermediate/dorsal | -0.8 | -3.23 | -3.26 |
| 40 | 18.2630333 | core | 1 | 0.83 | -5.8 | intermediate/dorsal | -0.6 | -3.18 | -3.37 |
| 30 | 6.61806667 | core | 0.8 | 1.29 | -5.71 | dorsal/dorsolateral | -1 | -3.09 | -3.16 |
| 41 | 23.3347667 | shell | 0.9 | -0.9 | -5.87 | intermediate/dorsal | -0.8 | 3.24 | -3.52 |
| 38 | 40.9856667 | core | 1.3 | -1 | -5.04 | dorsal/dorsolateral | -1.2 | 2.86 | -3.1 |
| 42 | 27.6835667 | shell | 1.1 | -0.83 | -5.85 | intermediate/dorsal | -0.9 | 3.21 | -3.43 |
| 29 | 58.1473333 | shell | 1.1 | 0.79 | -6.08 | intermediate/dorsal | -1 | -3.05 | -3.67 |

| | | | | | | | | | |
|---|---|---|---|---|---|---|---|---|---|
| 32 | 42.087 | core | 1.4 | 1.22 | -5.26 | intermediate/dorsal | -0.8 | -3.06 | -3.45 |
| 12 | 42.832 | shell | 1 | -0.44 | -5.67 | other | -0.7 | 3.4 | -3.03 |
| 70 | 73.8186667 | core | 1.2 | 1.02 | -5.68 | intermediate/dorsal | -1 | -3.33 | -3.35 |
| 55 | 40.7856667 | core | 1 | 0.96 | -5.69 | intermediate/dorsal | -1.2 | -3.11 | -3.53 |
| 60 | 64.743 | shell | 0.9 | -0.9 | -5.89 | intermediate/dorsal | -1.1 | 3.2 | -3.44 |
| 54 | 50.5396667 | core | 1.3 | 1.14 | -5.32 | dorsal/dorsolateral | -1.1 | -3.12 | -2.88 |
| 71 | 43.466 | core | 1.1 | 0.93 | -5.71 | intermediate/dorsal | -1 | -3.18 | -3.67 |
| 13 | 25.3479 | core | 0.9 | -0.85 | -5.59 | dorsal/dorsolateral | -1 | 3.05 | -2.87 |
| 11 | 20.8434 | core | 1.3 | -1.21 | -5.57 | intermediate/dorsal | -0.7 | 3.03 | -3.54 |
| 392 | 53.8986667 | other | 0.6 | 0.89 | -6.93 | intermediate | -1.5 | -3.38 | -3.38 |
| 61 | 43.5993333 | core | 0.9 | -1.01 | -5.84 | intermediate/dorsal | -0.9 | 3.31 | -3.42 |
| 10 | 42.3093333 | core | 1 | -0.99 | -5.56 | intermediate/dorsal | -0.9 | 3.13 | -3.44 |
| 312 | 67.49 | core | 1.1 | -1 | -5.9 | intermediate/dorsal | -0.7 | 2.99 | -3.74 |
| 313 | 46.38 | other | 0.3 | 1.06 | -6.33 | intermediate/dorsal | -0.8 | -3.08 | -3.49 |
| 314 | 43.8553333 | core | 1.1 | -0.96 | -5.89 | intermediate/dorsal | -0.9 | 3.24 | -3.44 |
| 302 | 62.2293333 | core | 0.9 | -1.28 | -5.67 | dorsal/dorsolateral | -1.2 | 3.14 | -3.04 |
| 303 | 58.3586667 | core | 0.6 | 0.99 | -5.53 | intermediate/dorsal | -0.7 | -3.58 | -3.54 |
| 301 | 52.2526667 | core | 1.4 | 1.18 | -4.67 | intermediate/dorsal | -0.8 | -3.22 | -3.29 |
| 375 | 14.4761333 | core | 1.1 | -0.89 | -5.66 | dorsal/dorsolateral | -0.7 | 3.1 | -2.95 |
| 600 | 63.3303333 | core | 1.2 | -1.15 | -5.12 | other | -0.6 | 2.97 | -2.4 |
| 363 | 43.5106667 | shell | 1.4 | 0.99 | -6.12 | intermediate/dorsal | -0.4 | -3.06 | -3.1 |
| 364 | 49.7166667 | shell | 1.4 | 0.76 | -6.12 | intermediate/dorsal | -0.7 | -3.21 | -3.35 |
| 376 | 55.945 | core | 1.2 | 1.01 | -5.77 | intermediate/dorsal | -0.8 | -3.23 | -3.49 |
| 58 | 46.3133333 | core | 1 | 1.32 | -5.79 | intermediate/dorsal | -0.7 | -3.3 | -3.2 |
| 361 | 54.299 | core | 1.3 | 1.01 | -5.48 | intermediate/dorsal | -0.6 | -3.03 | -3.02 |
| 360 | 52.7753333 | shell | 0.9 | -0.8 | -6.04 | other | -1.5 | -3.97 | -3.52 |
| 377 | 47.9936667 | core | 1.1 | -1.02 | -5.79 | intermediate/dorsal | -1.1 | 3.18 | -3.36 |
| 59 | 44.1223333 | core | 1 | -1.21 | -5.4 | intermediate/dorsal | -1 | 3.02 | -3.54 |
